# Archaeal histone HTkC hypercompacts DNA

**DOI:** 10.64898/2025.12.06.692711

**Authors:** Samuel Schwab, Yimin Hu, Shingo Yamamoto, Kodai Yamaura, Naomichi Takemata, Marcus D. Hartmann, Marc K.M. Cajili, Bert van Erp, Haruyuki Atomi, Vikram Alva, Birte Hernandez Alvarez, Remus T. Dame

## Abstract

Histones are important organizers of chromatin in eukaryotes and archaea. In eukaryotes, the core histones assemble with DNA to form the octameric nucleosome. In archaea, histones form hypernucleosomes that are not restricted to an octameric histone core but can extend to variable lengths. We previously identified face-to-face (FtF) histones as a widely distributed group of archaeal histones that assemble into toroidal tetramer structures, distinct from nucleosomal histones. Here, we characterize the FtF histone HTkC from *Thermococcus kodakarensis*, which also encodes the canonical histones HTkA and HTkB. We show that HTkC wraps DNA around its toroidal tetramer and forms highly compact nucleoprotein complexes, achieving a level of compaction approximately twice that of hypernucleosomes. Consistent with a major chromatin-organizing role, *htkC* is among the most highly expressed genes in *T. kodakarensis* and its deletion leads to impaired growth. Together, these findings establish FtF histones as important organizers of archaeal chromatin alongside classical histones.

## Introduction

Histones play an essential role in the chromatin organization of eukaryotes [1, 2]. In eukaryotes, histones H2A, H2B, H3, and H4 form, together with DNA, the nucleosome [3]. Despite the strong association of nucleosomes with the complex multicellular lifestyle of eukaryotes, nucleosomal histones are a prokaryotic invention. Most archaea encode histones that form hypernucleosomes, nucleosome-like structures that are not limited in size to an octameric histone core, but instead form nucleosomes of any size [4, 5, 6]. These archaeal nucleosomal histones were first discovered in 1990, and since then have been the focus of the prokaryotic histone field [7].

Despite forming complex nucleoprotein structures, histones are primarily defined by a small ∼70 amino acid-long histone fold. This fold includes three alpha helices, *α*1, *α*2, and *α*3, linked by two linkers, l1 and l2. These histone folds dimerize, with the *α*2-helices crossing and the *α*1 and *α*3 helices on opposite sides of the dimer. The dimer is the smallest functional unit and binds ∼30 base pairs without sequence specificity. In eukaryotes and Asgard archaea, histones commonly contain disordered N-terminal tails, which, in the case of eukaryotes, undergo extensive post-translational modifications (PTMs) that regulate nucleosome behavior [8]. The majority of prokaryotic histones, however, lack these tails and possess only the histone fold. Furthermore, while eukaryotic histones form obligate heterodimers, prokaryotic histones form homodimers and in some cases heterodimers, although little is known about the heterodimeric complexes [9].

The prokaryotic field has focused on histones HMfA and HMfB from *Methanothermus fervidus* and HTkA and HTkB from *Thermococcus kodakarensis* [4, 5, 6, 7, 10, 11, 12, 13, 14, 15, 16]. Recent bioinformatic studies, however, have described a wide range of new, uncharacterized histones in prokaryotes and viruses [17, 18]. Despite containing the histone fold, these histones are predicted to form different multimer structures compared to the canonical nucleosomal histones, allowing them to organize DNA in different ways. For example, the bacterial histone HBb from *Bdellovibrio bacteriovorus* does not form nucleosome-like structures, but rather functions as a histone dimer and bends DNA [19, 20]. New histones were also identified in the model organisms *M. fervidus* and *T. kodakarensis*. In *M. fervidus*, HMfC is capable of bridging DNA instead of wrapping DNA as a hypernucleosome [18]. Other DNA-bridging histones include MJ1647 from *Methanocaldococcus jannaschii* [21]. In *T. kodakarensis*, HTkC (TK1040) forms a toroidal tetramer in which two histone dimers face each other, unlike the V-shaped tetramer formed by canonical nucleosomal histones (Fig. 1) [18]. These face-to-face histones (FtF) are widely distributed across archaea and are also encoded by some bacteria. FtF histones are more closely related to the bacterial dimer histones, such as HBb, than to hypernucleosome-forming histones, and are part of a superfamily called the *α*3 histones, which are characterized by their short *α*3 helix.

**Figure 1:**
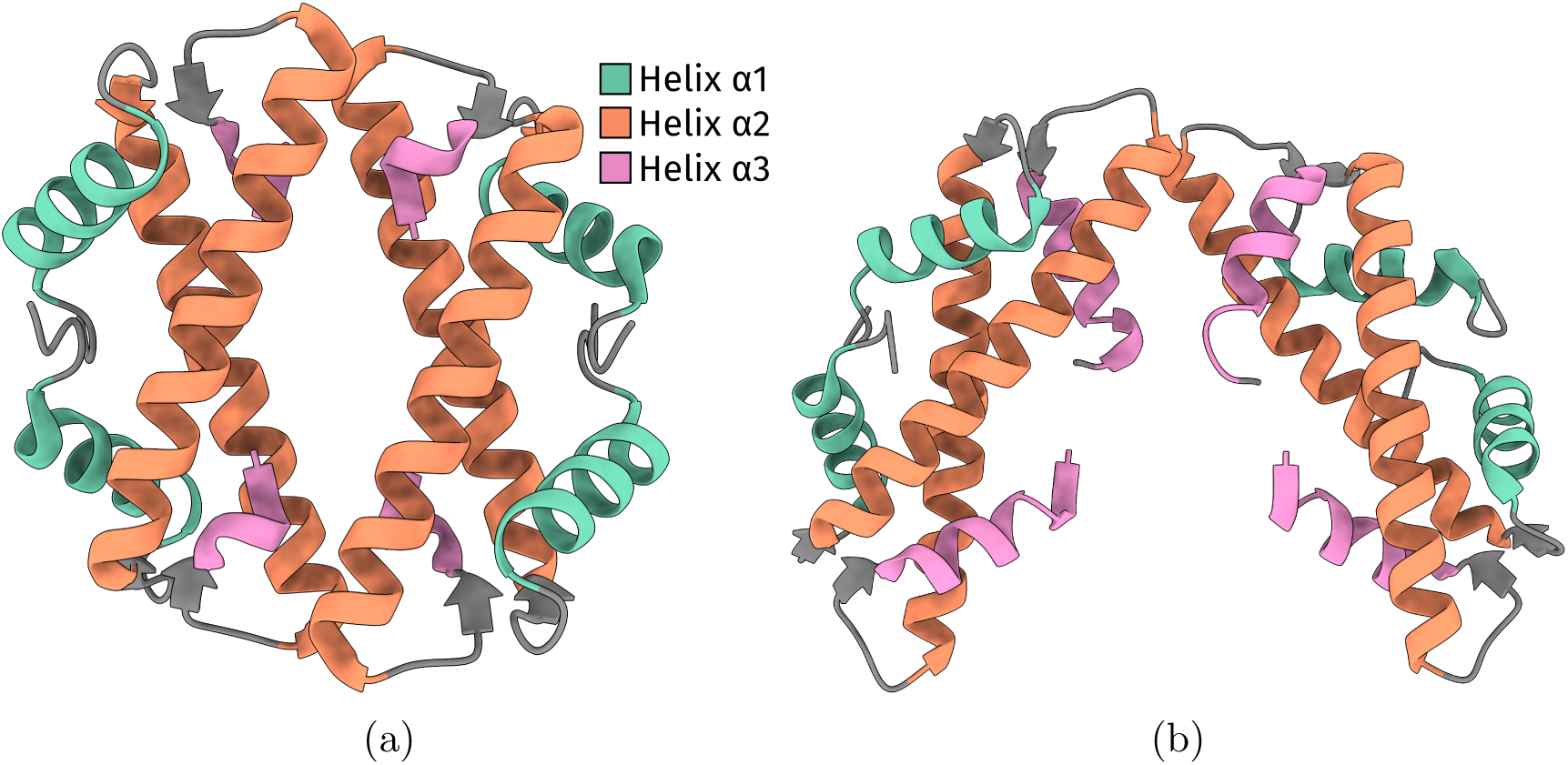
HTkC forms a tetramer structure distinct from canonical histones. (a) Crystal structure of HTkC (PDB: 9F2C) [18]. (b) Crystal structure of the canonical histone HMfB (PDB: 5T5K) [4].

FtF histones appear to be important chromatin organizers among prokaryotes. In the Leptospirales bacteria *Leptospira interrogans serovar Lai*, the FtF histone LA2458 is essential for cell viability [20]. In the archaeon *Haloferax volcanii*, the FtF histone is among the highest expressed genes in the organism [18]. The bacterial HLp from *Leptospira perolatii* is the only FtF histone to have been functionally characterized, demonstrating that it compacts DNA by wrapping it around its tetramer [22]. Apart from a crystal structure, little is known about the archaeal FtF histones and how they organize DNA [18].

In this study, we provide the first functional characterization of an archaeal FtF histone, HTkC from *T. kodakarensis*. Our findings demonstrate that HTkC is capable of wrapping DNA, forming extremely compact nucleoprotein structures, with a compaction level roughly twice that of hypernucleosomes. Furthermore, we show that *htkC* is highly expressed in *T. kodakarensis* and that its deletion impairs growth.

## Results

### Arginine 45 is critical for tetramer formation

We previously crystallized HTkC, revealing its toroidal tetramer structure (Fig. 1a) [18]. The dimer-dimer interface of the tetramer is positioned similarly to that of the hypernucleosome histone HMfB (Fig. 1b) [4]. However, in the HTkC tetramer, both of these interfaces are engaged within the closed toroid. As a result, no additional histone dimers can bind to these sites, preventing HTkC from forming an extended nucleosome-like structure. In the crystal structure, the HTkC tetramer is stabilized by arginine 45, which forms salt bridges with the C-terminal carboxyl group of the opposing dimer across the dimer-dimer interface (Fig. 2a and Supplementary Fig. 1). Arginine 45 is strongly conserved among FtF histones, suggesting that this residue is critical for tetramer formation [18]. To test whether arginine 45 is required for tetramer assembly, we overexpressed HTkC and HTkC R45A recombinantly in *Escherichia coli* and purified both proteins. We measured their oligomeric states using size-exclusion chromatography coupled with multi-angle static light scattering (SEC-MALS) (Fig. 2b and Supplementary Table 1). In SEC-MALS, wild-type HTkC elutes as a single peak corresponding to the tetrameric state. In contrast, HTkC R45A elutes as a dimer, confirming that arginine 45 is critical for tetramer formation. To evaluate the impact of the R45A substitution on protein stability, we performed far-UV circular dichroism (CD) spectroscopy across a temperature gradient. Wildtype HTkC retained its characteristic *α*-helical spectrum with only modest changes up to 95 ^◦^C (Fig. 2c), reflecting its intrinsic thermostability and aligning with the thermophilic lifestyle of the host organism *Thermococcus kodakarensis*. The R45A mutant, however, showed an earlier and more pronounced decrease in ellipticity—particularly at 222nm—indicating reduced thermal stability and partial unfolding at 95 ^◦^C (Fig. 2d).

**Figure 2:**
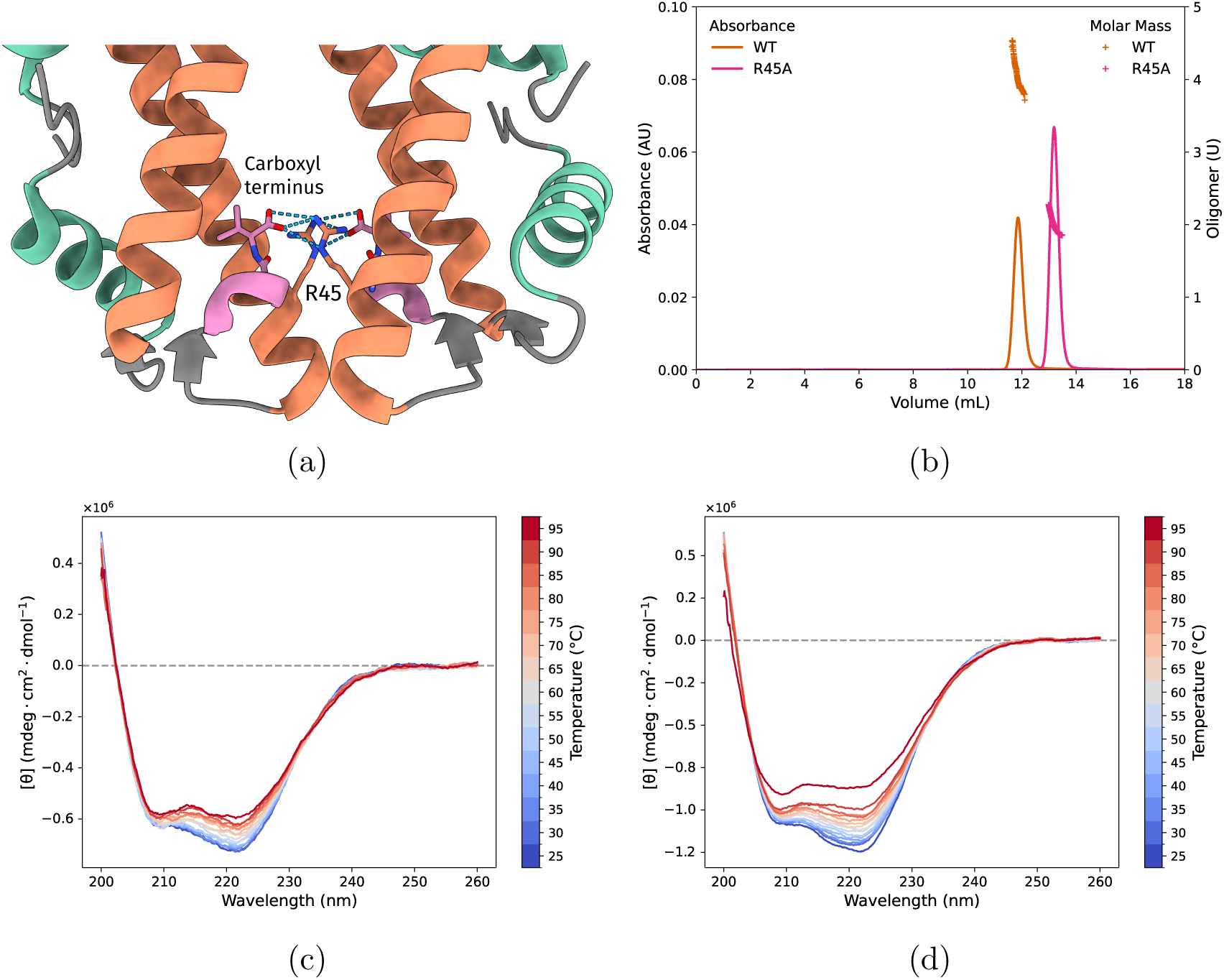
Arginine 45 is necessary for tetramerization of HTkC. (a) Closeup of the interactions between arginine 45 and the C-terminal carboxyl group (PDB: 9F2C) [18]. Helices *α*2 and *α*3 are colored orange and pink, respectively. (b) UV absorbance and molar mass profiles as measured by SEC-MALS for wildtype HTkC (orange) and HTkC R45A (pink). Molar mass is expressed as the number of subunits in the oligomer. (c,d) CD spectra for (c) wild-type HTkC and (d) HTkC R45A at temperatures ranging from 25 ^◦^C to 95 ^◦^C.

### HTkC wraps and (hyper)compacts DNA in vitro

We previously hypothesized that HTkC might wrap ∼50 bp of DNA around its tetramer [18]. To investigate whether HTkC binds to DNA, we performed electrophoretic mobility shift assays (EMSAs) using a 150 bp DNA fragment. HTkC readily binds the DNA fragment, resulting in two distinct shifted species corresponding to two bound states (Fig. 3a). From the EMSA alone, it is not clear whether these two states reflect different stoichiometries of HTkC or distinct nucleoprotein structures. At higher concentrations of HTkC, a larger, aggregate-like, complex appears. In contrast to the wild-type, the R45A mutant does not form distinct bound states (Fig. 3b). Instead, the nucleoprotein complex appears as a smeared band. This EMSA behavior is similar to that of the bacterial DNAbending histone HBb and the DNA-bending nucleoid-associated proteins (NAPs) such as Cren7 [19, 23], suggesting that HTkC R45A functions as a dimeric DNAbending histone.

**Figure 3:**
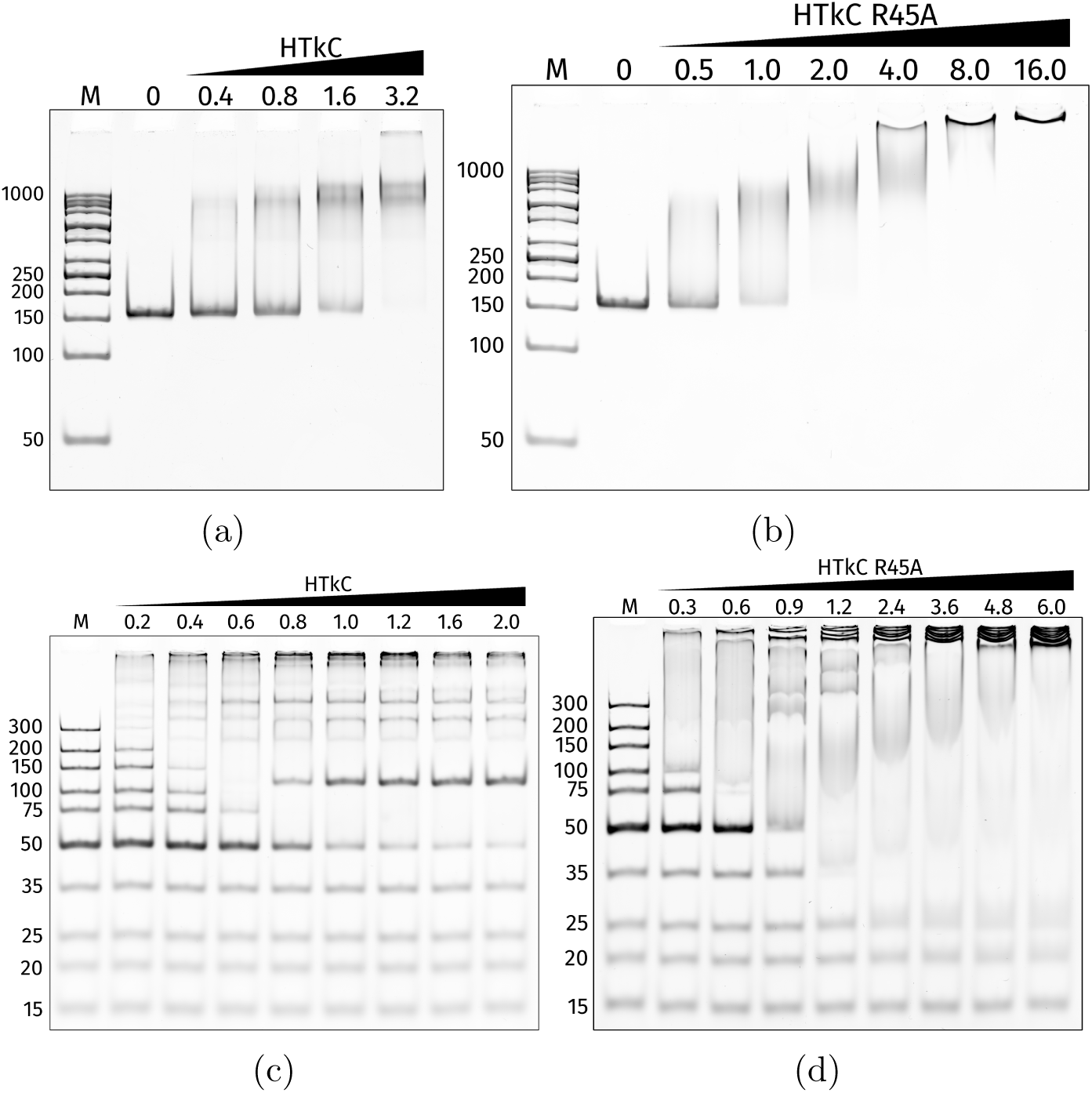
HTkC binds to DNA fragments. ≥**50 bp.** (a,b) EMSAs of (a) wildtype HTkC and (b) HTkC R45A in the presence of the 150 bp DNA fragment. The protein:DNA mass ratios are indicated above the lanes. (c,d) EMSAs of (c) wild-type HTkC and (d) HTkC R45A in the presence of the GeneRuler Ultra Low Range DNA Ladder (Thermo Fisher Scientific). The protein:DNA mass ratios are indicated above the lanes.

To identify the smallest DNA-fragment that HTkC can bind, we performed EMSAs with a DNA-ladder. HTkC binds all DNA fragments down to 50 bp (Fig. 3c). In contrast, the R45A mutant binds smaller fragments with clear shifts observed down to 35 bp (Fig. 3d). The 25 bp and 15 bp fragments do not produce distinct shifted bands but appear blurry, suggesting transient or unstable interaction of HTkC R45A with these short fragments. The difference in minimal DNA-binding length between wild-type and R45A suggests that the two histone dimers within the tetramer do not bind DNA independently of each other. Such a mode of binding would enable the tetramer to bridge two separate DNA duplexes. However, a DNA-bridging assay, which we previously used to measure bridging by histones HMfC and MJ1647 [18, 21] showed no bridging activity for HTkC (Supplementary Fig. 3a).

To further investigate the nucleoprotein complexes formed by HTkC, we performed a ligase-mediated circularization assay to test for DNA-bending and wrapping. In this assay, HTkC is incubated with a 240 bp DNA fragment. Subsequently, a ligation reaction is started by the addition of T4 ligase. If HTkC bends or wraps DNA, this will promote the formation of DNA circles. Small circles are unambiguously identified after digestion with Exonuclease V, followed by separation through gel electrophoresis. With increasing concentrations of both wild-type HTkC and HTkC R45A, we observe the formation of small circles, confirming that both proteins bend or wrap DNA (Fig. 4a and 4b).

**Figure 4:**
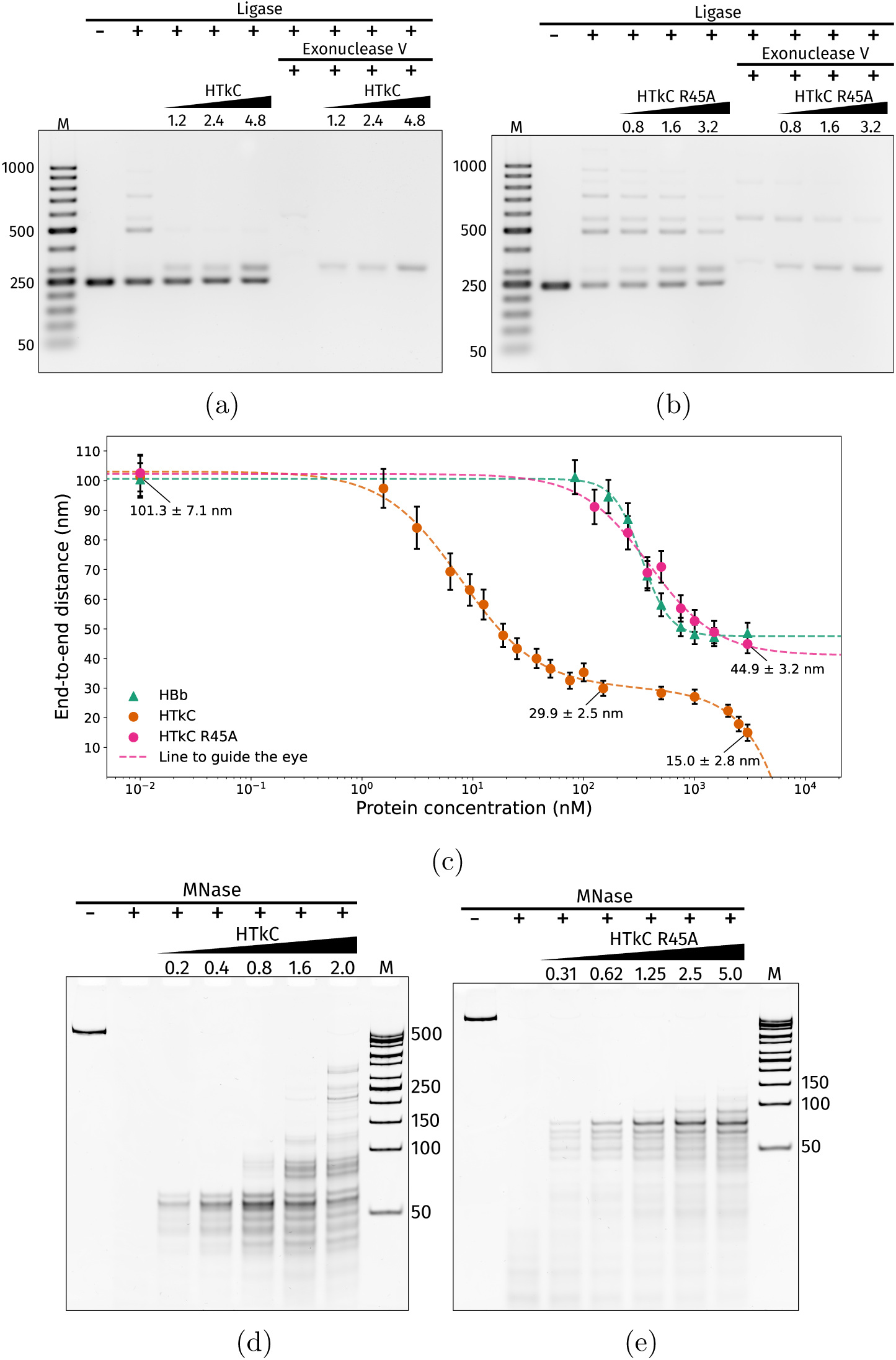
(Caption is on the next page.) Figure 4: **HTkC wraps and (hyper)compacts DNA.** (a, b) Ligase-mediated circularization assay of 240 bp DNA with (a) HTkC and (b) HTkC R45A. ProteinDNA mass ratios are indicated above the lanes. DNA circles are identified by Exonuclease V treatment. (c) TPM experiment of HTkC and HTkC R45A with 685 bp DNA. TPM data from HBb are shown as a reference [19]. All measurement points represent averages of triplicate measurements, with error bars indicating standard deviations. (d, e) In vitro MNase digestion assay of (d) HTkC and (e) HTkC R45A with 685 bp DNA. Protein:DNA mass ratios are indicated above the lanes.

To quantify DNA compaction by HTkC, we performed tethered particle motion (TPM) analysis, a single-molecule technique that allows real-time observation of DNA-protein interactions. In TPM, a linear (685 bp) DNA molecule is tethered to a glass surface and to a polystyrene bead at opposite ends. The position of the bead is tracked over time using a light microscope. From the ensemble of positions, we calculate the root-mean-square (RMS) displacement of the bead (Supplementary Fig. 3b) and the end-to-end (EtE) distance of the DNA (Fig. 4c). Proteins that compact DNA reduce the EtE distance, which is observed in TPM as a reduction of the RMS displacement of the bead. With increasing concentrations of HTkC, we observe two distinct regimes of compaction (Fig. 4c). In the first regime (0 to 150 nM), HTkC compacts the DNA down to an EtE distance of 29.9 ± 2.5nm.

This level of compaction is similar to that observed for archaeal hypernucleosomes formed by HMfA, HMfB, HTkA, and HTkB, suggesting that HTkC wraps DNA [5, 16]. In the range of 150 nM to 500 nM, the EtE distance remains constant at ∼30nm. From 500 nM to 3000 nM, we observe a second regime of compaction, with the EtE distance decreasing to 15.0 ± 2.8nm. The EtE distance is therefore halved compared to the low concentration regime, indicating that this state is significantly more compact. We are unable to measure at concentrations above 3000 nM as the movement of the bead becomes too small to observe with our light microscope. The R45A mutant lacks the two-regime phenotype and shows significantly less compaction compared to the wild-type (Fig. 4c). The EtE distance is reduced to 44.9 ± 3.2nm, a value that is characteristic for DNA-bending proteins, such as HU, Cren7, and histone HBb [19, 24, 25]. HBb exhibits a highly similar TPM profile, suggesting that HTkC R45A bends DNA as a dimer, in the same manner as HBb.

To investigated whether the HTkC tetramers form larger protein-DNA complexes, we performed a micrococcal nuclease (MNase) digestion assay. In the MNase assay, HTkC is incubated with linear DNA. MNase is subsequently added to digest all DNA that has not been protected by protein. For wild-type HTkC, we observe protected fragments that are multiples of 30 bp (60, 90, 120, etc.), similar to MNase assays of hypernucleosomes (Fig. 4d) [6, 19]. This suggests the presence of a larger ordered nucleoprotein complex with the HTkC tetramers tightly packed on the DNA. The ladder pattern is lost in HTkC R45A, indicating that tetramer formation is crucial for forming the larger nucleoprotein complexes (Fig. 4e).

### DNA encircles around the HTkC tetramer

HTkC forms larger nucleoprotein structures, as indicated by the in vitro MNase and TPM experiments. However, the architecture of these complexes remains unclear. To determine the structure of HTkC bound to DNA, we co-crystallized HTkC in the presence of a 30-bp double-stranded DNA (dsDNA) fragment. HTkC successfully crystallized within a week. Diffraction data for the HTkC-DNA complex were collected and processed to a resolution of 1.95 Å, and the structure was determined by molecular replacement using the DNA-free HTkC coordinates (PDB: 9F2C [18]) as a search model. The asymmetric unit contains one HTkC dimer bound to a 29-nucleotide DNA strand. Via crystallographic symmetry, these strands assemble into a seemingly continuous DNA duplex winding through the crystal lattice, encircling half of each HTkC tetramer (see Methods; Fig. 5a-c and Supplementary Fig. 4). To assess whether the lattice arrangement corresponds to either the low- or high-concentration regime in TPM, we measured the EtE distance of a 685-bp DNA fragment in the lattice. This EtE distance (∼103 nm) is nearly an order of magnitude larger than the compacted state measured in the low-concentration regime in TPM. Thus, the crystal lattice does not represent the compacted structures that HTkC forms in solution, as measured by TPM. Nevertheless, the crystal structure provides insights into how individual HTkC tetramers interact with DNA. The crystal structure reveals that the HTkC tetramer engages the DNA minor groove through residues in the N-terminal region of the *α*1-helices and loops l1 and l2. In the asymmetric unit, key contacts involve residues K7, S8, K9, R51, K52, T53, and Y55 from one monomer, and K7, S8, K9, K11, R22, and V23 from the other (Fig. 5d and 5e). All of these residues interact with the phosphate backbone or ribose moieties of the DNA, consistent with non-sequencespecific binding. Overall, HTkC exhibits similarities to both the nucleosomal histone HMfB and the bacterial dimer histone HBb. S8, K9, K11, K52, and T53 are DNA-binding residues in the related bacterial histones HBb and HLp [19, 22], while K11, R22, K52, and T53 are known DNA-binding residues in HMfB [4]. Like in HMfB, R22 inserts itself into the minor groove and is the only residue that approaches nucleobases. HTkC also contains a unique DNA-binding residue not found in other histones. This residue, R51, lies within the RxTxxxxD motif, a strongly conserved feature in histones from eukaryotes, archaea, and bacteria that positions the l2 loop and helix *α*3 via a conserved arginine-aspartate contact (Supplementary Fig. 1). R51 is therefore normally not involved in DNA-interactions. However, in HTkC, the aspartate in this motif is replaced by a histidine (Supplementary Fig. 2). As a result, R51 points outward and contacts the DNA. To maintain the position of the l2 loop and helix *α*3, HTkC instead forms salt bridges between E49, R57, and H58.

**Figure 5:**
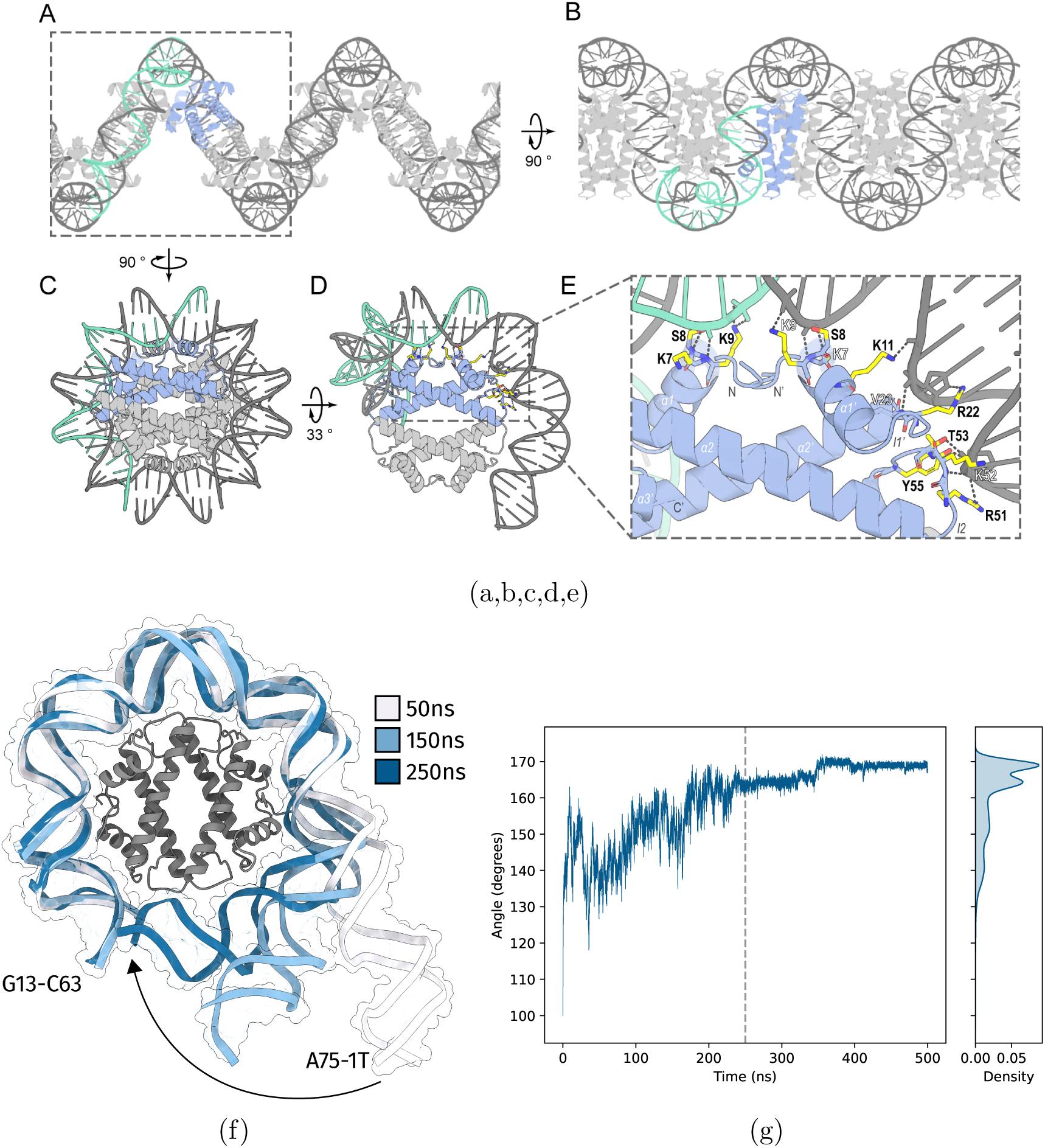
Structure and molecular dynamics simulations of HTkC with DNA. (a-c) Crystal packing of the HTkC-DNA complex viewed from different angles, with selected symmetry mates generated within 4 Å. (d) HTkC tetramer wrapped by dsDNA (PDB: 9T30) is shown in cartoon representation with DNAbinding residues shown as sticks. (e) Close-up of residues involved in DNA-binding. In all panels, the contents of a single asymmetric unit are shown in colors, selected symmetry mates in gray, and hydrogen bonds and salt bridges are indicated as dashed lines. (f) Three frames from simulation #1 at 50, 150, and 250 ns that show the trajectory of wrapping. Base pairs C1-G75 to C12-G64 are not shown to reduce visual clutter. (g) The bending angle in degrees between base pairs G13C63, A75-T1, and the middle base pair (C30-G46) during simulation #1. The gray dashed line highlights the time point at which complete wrapping occurs (250ns). See Supplementary Fig. 6 for the MD data from simulation #2.

The co-crystal structure shows that the HTkC tetramer wraps DNA halfway around itself. Owing to the dihedral D_2_ symmetry of the tetramer, we expect that HTkC could wrap DNA fully around the tetramer. To examine this possibility, we performed molecular dynamics (MD) simulations on the co-crystal structure (see Data availability; Supplementary Fig. 5 and 6). As the starting model, we used a single tetramer with 75 bp of DNA taken from the crystal lattice (Supplementary Fig. 5a). We performed two simulations of 500ns each (1µs total). During the first 250ns of simulation #1, the DNA progressively wraps around the whole tetramer (Fig. 5f). We measured the state of wrapping along the simulation by calculating the “bending” angle formed between base pairs G13-C63, A75-T1, which correspond to the outermost base pairs of the fully wrapped state, and the midpoint between them (Fig. 5g) (see Methods). The bending angle increases over the first 250ns, consistent with base pairs G13-C63 and A75-T1 being bent closer to each other. The number of protein-DNA contacts also increases steadily over this interval (Supplementary Fig. 5b). At 250ns, the bending angle reaches a stable value of ∼170^◦^, near the theoretical maximum of 180^◦^ (Fig. 5g). In the final 250ns, the DNA remains strongly bent into a mini-circle around the tetramer (Fig. 5f). The key residues that bind and bend the DNA are the same as those identified from the co-crystal structure (Supplementary Fig. 5c and 5d). Simulation #2 shows a similar progression: the tetramer fully wraps DNA within 150ns and remains wrapped for 200ns, after which one DNA arm partially unwraps from the tetramer. This unwrapping event is caused by steric hindrance between the two DNA ends. Because the HTkC tetramer is completely planar, the DNA ends collide after completing a full turn around the tetramer. This steric clash is expected to be more problematic for longer DNA fragments than for short ones such as the 75 bp fragment used in our simulations. How this steric hindrance is resolved when HTkC binds to longer DNA fragments remains unclear.

**Figure 6:**
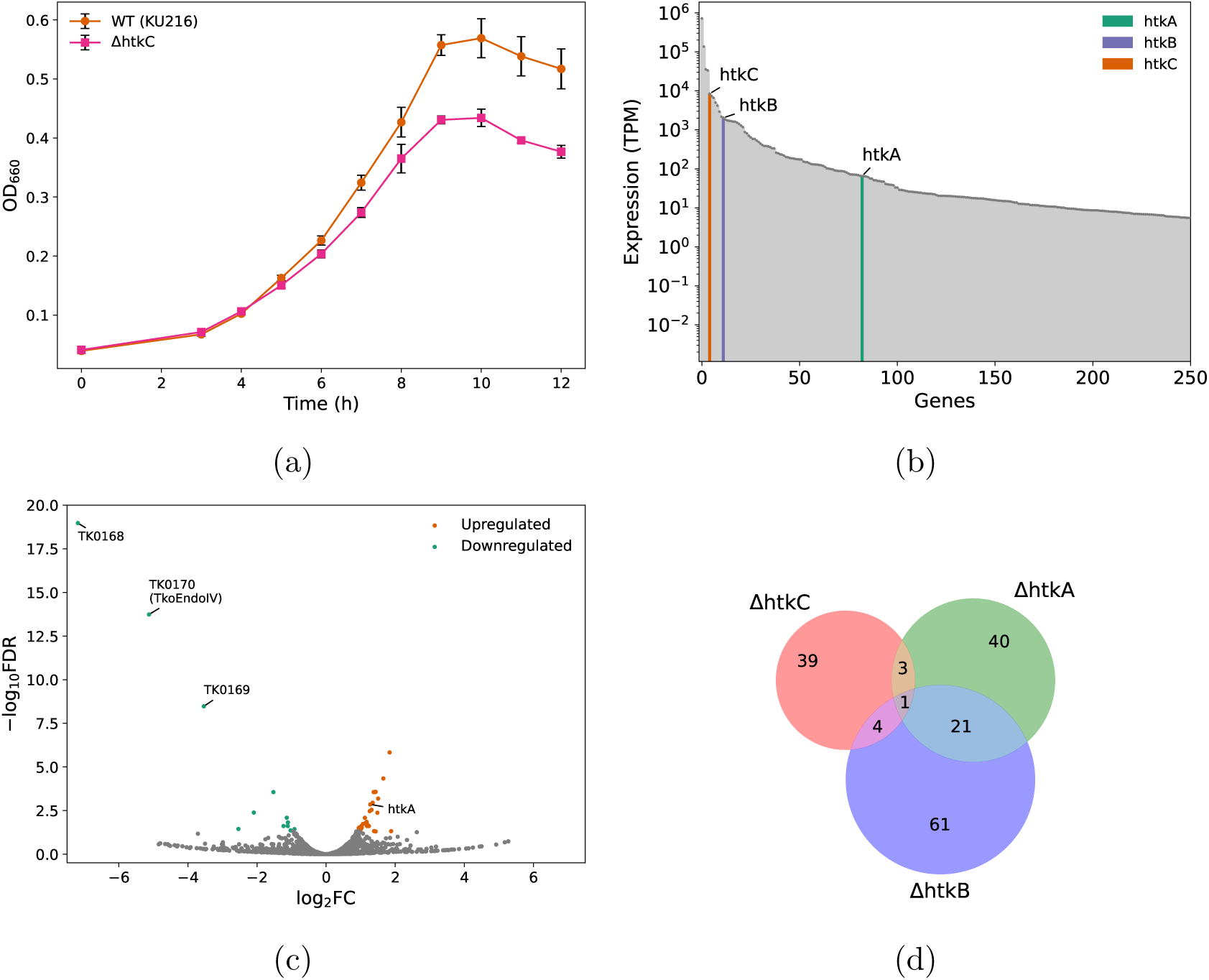
HTkC is highly expressed in. *T. kodakarensis* **and its deletion impairs growth.** (a) Growth curves of the *T. kodakarensis* strain KU216 and its derivative Δ*htkC* strain. (b) Expression levels, calculated as Transcripts per Kilobase Million (TPM), of the top 250 expressed genes from biological replicate 1. Genes are ordered from highest (left) to lowest (right) expression. The *htkA*, *htkB*, and *htkC* genes are highlighted in green, purple, and orange, respectively. For the other two biological replicates, see Supplementary Figure 8. (c) Volcano plot of transcriptomic changes in the Δ*htkC* strain compared to KU216. Orange and green dots indicate upregulated and downregulated genes, respectively, with an FDR ≤ 0.05. (d) Venn diagram of differentially expressed genes of the Δ*htkC*, Δ*htkA*, and Δ*htkB* strains [26].

We observe alternating widening and narrowing of the major and minor grooves in both MD simulations (Supplementary Fig. 7). Severe widening of the major groove occurs at the locations where HTkC engages the minor groove, with major grooves being up to 2.5 times wider than in normal B-DNA. At these same sites, the minor groove narrows to roughly half of its normal width. Between binding sites, the major groove is slightly narrowed and the minor slightly widened. These alterations in major and minor groove width are also observed in the related bacterial histone HBb and archaeal histone HMfB when bound to DNA (Supplementary Fig. 7c) [19].

### HTkC is highly expressed and affects growth

FtF histones, such as HTkC, are widely distributed throughout archaea and are also present in several bacteria, such as in the Leptospirales order [18]. In *Leptospira interrogans serovar Lai*, the FtF histone homolog is essential for cell viability [20, 22]. In Thermococci, FtF histones are not strictly conserved, suggesting a non-essential role [11]. We successfully deleted *htkC* from the *T. kodakarensis* strain KU216, confirming its non-essential role under laboratory conditions. When grown at 85 ^◦^C in nutrient-rich medium (ASW-YT-m1-S^0^), the Δ*htkC* strain exhibits reduced growth compared to the parental strain (Fig. 6a). Notably, in the Δ*htkC* strain, the cell density at the stationary phase is reduced by 27% relative to KU216. This growth deficiency is similar to that observed when histone HTkA is deleted, which results in a 25% reduction in stationary phase cell density [26].

To investigate the expression of *htkC* and transcriptomic changes upon its loss, we performed RNA-seq on the KU216 and Δ*htkC* strains. Transcriptomic data were obtained in triplicate for both strains in early log phase. In KU216, *htkC* is among the top 10 highest expressed genes, with expression levels 4-24 times higher than *htkB* and 129-421 times higher than *htkA* (Fig. 6b and Supplementary Fig. 8). In the Δ*htkC* strain, 45 significantly differentially expressed genes (DEGs; FDR *<* 0.05) were identified compared to KU216 (Fig. 6c). Out of these, 34 DEGs were upregulated. Importantly, *htkA* shows a 2-fold increase in expression, suggesting that the loss of *htkC* may be compensated for by increased *htkA* expression. For other chromatin-related proteins, including SMC (TK1017), ScpA (TK1018), ScpB (TK1962), TrmBL2 (TK0471), HTkB (TK2289), and the DUF1931 histone (TK0750), we detected no significant change in expression. Other upregulated genes include those encoding various hypothetical importers/exporters, ribosomal proteins, and the RNA polymerase subunit Rpo12 (See Data availability). Among the downregulated DEGs, 3 gene loci are strongly downregulated (fold changes ≥ 3): TK0168, TK0169, and TK0170. The gene TK0170 encodes TkoEndoIV, a major apurinic/apyrimidinic endonuclease involved in base excision repair and essential for cell viability [27]. The reduced growth observed in the Δ*htkC* strain may be partly due to the 35-fold decrease in TK0170 expression. The other two genes, TK0168 and TK0169, encode an AsnC/Lrp-like ligand binding domain and a MJ1563/MarR-like transcription factor, respectively, that together form a two-gene operon adjacent to TK0170. The transcriptional effects after deletion of *htkC* appear minor with only 45 DEGs. This is in line with the deletion of the other histones, *htkA* and *htkB*, which result in 65 and 87 DEGs, respectively, as measured by microarray hybridization [26]. 8 of the 45 DEGs in the Δ*htkC* strain are also DEGs in either Δ*htkA* or Δ*htkB* (Fig. 6d). However, all but one of these DEGs show opposite effects in the Δ*htkC* strain, i.e., upregulated genes are downregulated and vice versa.

## Discussion

In this study, we employ in vitro, in vivo, and computational methods to characterize the archaeal FtF histone HTkC from *T. kodakarensis*. We show that nucleosomal histones are not the only primary chromatin-organizing histones in archaea, but that FtF histones, which are widely distributed, are also important organizers of archaeal chromatin.

HTkC forms a toroidal tetramer structure, unlike the V-shaped tetramer of nucleosomal histones. HTkC wraps DNA around this tetramer core and is capable of forming larger nucleoprotein complexes. Based on our TPM results, HTkC forms two types of nucleoprotein complexes. At low concentrations, HTkC compacts DNA to a level similar to hypernucleosomes [5, 16]. At higher concentrations, the nucleoprotein complexes are at least two-fold more compact than hypernucleosomes. We are unable to reach saturation in the high concentration regime, because the complexes become too compact to measure in our TPM assay.

The structures adopted by the low- and high-concentration nucleoprotein complexes remain unclear. We attempted to co-crystallize HTkC and DNA, but the resulting crystal lattice is not compact enough to represent either the low- or high-concentration states. Strikingly, however, the DNA in the crystal is bent into a helix, similar to hypernucleosomes [4]. A structure in which HTkC tetramers stack upon each other to form a histone core around which the DNA is wrapped would provide compaction levels similar to hypernucleosomes, and could therefore correspond to the structure observed at low concentrations. Hypothesizing possible structures for the high-concentration complexes is more difficult, as HTkC is the only protein for which we have observed such levels of compaction in our TPM assay. One possibility is a fiber-like arrangement, reminiscent of the 2-start nucleosome fiber observed for eukaryotic histones [28], as such fibers provide substantially more compaction than a hypernucleosome.

FtF histones are one of the few histone types that are encoded by both archaea and bacteria. The only other characterized FtF histone is the bacterial histone HLp from *L. perolatii*, which shows low sequence identity with HTkC (29%) [22]. Both HLp and HTkC wrap DNA around their tetramers, and given that all FtF histones are predicted to form toroidal tetramer structures, this binding mode is likely conserved among FtF histones [18]. Key differences between HLp and HTkC include the absence of a high-concentration compaction state in TPM for HLp and the lack of a well-defined 30-bp protection pattern in the MNase assay. Together, these suggest that the higher-order complexes formed by HTkC on DNA differ from those formed by HLp, highlighting an important difference between archaeal and bacterial FtF histones. Nevertheless, FtF histones appear to be principal DNA-organizing proteins in both archaea and bacteria

The behavior of the HTkC R45A mutant illustrates how a single conserved residue in the FtF scaffold can alter the DNA-binding mode. The R45A mutant is highly reminiscent in its phenotype of the bacterial dimer histone HBb, suggesting that it simply bends DNA as a dimer rather than wrapping it as a tetramer. Both HBb and HTkC are part of the *α*3 histone superfamily. In the case of HTkC, a truncated *α*3 helix is likely critical, as extension of the C-terminus will disrupt the salt bridges between R45 and C-terminal carboxyl group. Among other FtF histones, R45 is one of the strongest conserved residues, although lysine is occasionally observed at this position [18]. Lysine should also be able to promote tetramerization, albeit with weaker interactions than arginine. Even though HBb is from a different domain of life, HTkC and HBb share higher sequence identity (42%) than HTkC does with HTkA/HTkB (26%). HBb, HTkC, and other *α*3 histones likely derive from a common ancestor. The lineage in which this ancestor originated is not entirely clear. FtF and nucleosomal histones, or an ancestor of both, were likely already present in the last common archaeal ancestor, as both FtF and nucleosomal histones are widely encoded within archaea. However, some bacteria also encode FtF (and bacterial dimer) histones. This either implies that FtF histones have been exchanged between the two domains through horizontal gene transfer or that the histone ancestor can be traced back to the last common ancestor of archaea and bacteria. In the former scenario, an FtF histone encoded by a bacterium may have lost its R45 at some point, giving rise to modern bacterial dimer histones.

FtF histones are widely encoded in archaea, including Euryarchaea, Asgard archaea, DPANN, TACK, and Thermoplasmatota [18]. Like the bacterial FtF and archaeal hypernucleosome histones, HTkC appears to be an important factor in chromatin organization, as illustrated by the reduced growth of the Δ*htkC* strain. Importantly, HTkC is the most highly expressed histone in early log phase in *T. kodakarensis*. This is supported by RNA-seq data obtained independently by Yamaura et al. (2025) [29]. However, in the RNA-seq data from Jäger et al. (2014), HTkC is the lowest expressed histone [30]. We hypothesize that the expression of *htkA*, *htkB*, and *htkC* may vary depending on growth conditions. Indeed, in the related organism *Thermococcus onnurineus* NA1, the relative expression levels of the hypernucleosome histones and the FtF histone change depending on the growth medium, with the FtF histone being the most highly expressed histone and among the top 5% of expressed genes in MM1-Formate medium [31]. An important open question concerns the functional relationship between HTkA, HTkB, and HTkC: does HTkC interact with the other two histones, structurally or functionally? The upregulation of *htkA* in the Δ*htkC* strain implies a functional relationship between the two. However, their functional roles are not fully redundant, as a double knockout of *htkA* and *htkB* is lethal, indicating that HTkC cannot compensate for the loss of HTkA and HTkB [26]. Several archaea, such as *Haloferax volcanii* and *Heimdallarchaeota LC2*, encode FtF histones but lack nucleosomal histones [18], suggesting that the function of FtF histones in archaea is not exclusively tied to nucleosomal histones.

In summary, our work establishes FtF histones as a new major group of DNA-wrapping histones that organize chromatin in archaea. Future studies will be needed to elucidate the molecular details of the compact nucleoprotein structures formed by HTkC and how or whether HTkC interacts with HTkA and HTkB.

Furthermore, characterization of other archaeal FtF histones will be important to reveal conserved and divergent properties within this group.

## Methods

### Expression, and purification of HTkC

HTkC (TK1040) was expressed and purified as previously described [18]. Briefly, pRD503 (Addgene: 220788) was transformed into chemically competent BL21(DE3)pLysS cells (Novagen). The culture was grown at 37 ^◦^C with 200RPM shaking, and upon reaching an OD600 of approximately 0.6, IPTG was added to a final concentration of 1mM to induce expression. Cells were harvested after 1 hour of induction and lysis was done with a Stansted Pressure Cell Homogenizer S-PCH-10 (Homogenising Systems Ltd.). The resulting supernatant was then heated at 70 ^◦^C for 15 minutes and purified with a HiTrap Heparin HP (Cytiva) column and a Superdex 75 Increase 10/300 GL size exclusion column.

### Cloning, expression, and purification of HTkC mutant R45A

The R45A mutation was introduced by PCR amplification of *htkC* from pRD503 with the R45A mutation present in the primers. The amplified product was subsequently cloned into a pET30b expression vector through Gibson Assembly. The sequence of the construct was verified by DNA Sanger sequencing (BaseClear). The plasmid (pRD661) was deposited at Addgene (Supplementary Table 2). The sequences of the oligonucleotides are available in the Supplementary Information (Supplementary Table 4).

pRD661 was transformed into chemically competent BL21(DE3)pLysS cells (Novagen) through heat shock. Lysogenic broth (LB) supplemented with 50µgmL^−1^ kanamycin and 25µgmL^−1^ chloramphenicol was inoculated with 1% (v/v) of a starter culture of transformed cells grown overnight. The culture was grown at 37 ^◦^C with 200RPM shaking, and upon reaching an OD600 of approximately 0.6, IPTG was added to a final concentration of 1mM to induce expression. After growing the cells further for 1 hour, the culture was harvested at 7510xg for 30 minutes at 4 ^◦^C. The cell pellet was resuspended in 50mM Tris-HCl, pH 8, 150mM NaCl, 10mM MgCl_2_, and stored at −70 ^◦^C. After thawing the cell pellet, PMSF and DNase were added, and lysis was done with a Stansted Pressure Cell Homogenizer S-PCH-10 (Homogenising Systems Ltd.) at 310MPa. The lysate was centrifuged at 16000xg for 30 minutes at 4 ^◦^C. The resulting supernatant was then heated at 70 ^◦^C for 15 minutes, followed by another round of centrifugation at 16000xg for 30 minutes at 4 ^◦^C.

The supernatant was run on a pre-equilibrated 5mL HiTrap Heparin HP (Cytiva) attached to an NGC Chromatography System (BioRad). A 50mL gradient of 50mM Tris-HCl, pH 8, 100mM KCl to 50mM Tris-HCl, pH 8, 1M KCl was applied, and the eluent was fractionated into 1mL fractions. After SDS-PAGE verification, the fractions containing the protein were pooled and concentrated using Vivaspin^®^ Turbo 4 3000 MWCO centrifugal concentrators. The concentrated sample was applied to a Superdex 75 Increase 10/300 GL size exclusion column equilibrated with 100mM KCl, 50mM Tris-HCl, pH 7.0, 10% glycerol. The protein present in the collected fractions corresponding to the peak was then characterized with SDS-PAGE and intact protein LC-MS and stored at −70 ^◦^C.

### Size exclusion chromatography coupled with multi-angle light scattering (SEC-MALS)

SEC-MALS analysis was performed using a system containing a miniDAWN TREOS, DynaPro NanoStar, Opti-lab differential refractometer (Wyatt technology), and 1260 Infinity II multiple wavelength absorbance detector (Agilent). Samples were run on a Superdex Increase 75 10/300 column (Cytiva) with 200mM NaCl, 50mM Tris, pH 7.0 as the running buffer at a flow rate of 0.75mLmin^−1^. Data was processed using the ASTRA 8 software package and plotted in Python with the Matplotlib package (see Code availability) [32, 33].

### Circular dichroism (CD) spectroscopy

Measurements were performed using a Jasco J-1500 CD Spectrometer in a 0.1mm quartz cuvette. Spectra were measured from 260nm to 200nm with 100ngµL^−1^ protein in a 10mM Potassium Phosphate (pH 7.0), 75mM KCl buffer. From 25 ^◦^C to 95 ^◦^C, spectra were recorded at a 5 ^◦^C interval with a 1 ^◦^Cmin^−1^ slope, scanning speed of 100nmmin^−1^, 0.1nm data pitch, 1nm band width, 2s D.I.T., and 200mdeg*/*0.1OD CD scale. At each temperature, 3 spectra were recorded and averaged. Spectra were plotted in Python with the Matplotlib package (see Code availability) [32, 33].

### Electrophoretic mobility shift assay (EMSA)

To test for DNA-binding, a 150 bp DNA fragment was used. To approximate the length of DNA fragments required for protein binding, GeneRuler Ultra Low Range DNA Ladder (ThermoFisher Scientific) was used as a substrate. The 150 bp fragment was generated by PCR amplification of pRD121 [5]. The sequences of the oligonucleotides are available in the Supplementary Information (Supplementary Table 4).

The DNA substrates were mixed with either HTkC or HTkC R45A in 50mM Tris-HCl (pH 7.0), 75mM KCl, 14.5% glycerol at the indicated ratios and incubated at room temperature for 30min. Subsequently, the samples were separated on a 7% polyacrylamide (0.5x TBE) gel for the 150 bp fragment or a 10% polyacrylamide (0.5x TBE) gel for the ladder substrate at 120V and 4 ^◦^C with 0.5x TBE as the running buffer. Gels were stained with 1x GelRed (Biotium) and imaged using the InGenius 5 imaging system (Syngene).

### DNA-bridging assay

The DNA-bridging assay was performed using a 47% GC 685 bp DNA substrate as previously described [18, 21, 34]. In short, the DNA substrates were generated by PCR using Thermo Scientific^®^ Phusion^®^ High-Fidelity DNA Polymerase for prey DNA and with a biotinylated primer for bait DNA. The subsequent PCR products were purified using a GeneElute PCR Clean-up kit (Sigma Aldrich) and prey DNA was 32P labeled with Polynucleotide Kinase and *γ*-32P ATP. For each bridging assay measurement, 6µL of streptavidin-coated paramagnetic beads were washed with 50µL PBS (12mM NaPO4 pH 7.4, 137mM NaCl). The beads were subsequently washed twice with 50µL CB (20mM Tris-HCl pH 8.0, 2mM EDTA, 2M NaCl, 2mg mL^−1^ Acetylated BSA, 0.04% Tween20) and then resuspended in 6µL of CB. Next, 100pM (in a total volume of 3µL) of biotinylated bait DNA was added to half of the suspension and incubated with the beads at 25 °C for 20 min in an Eppendorf Thermomixer with an Eppendorf Smartblock 1.5 mL at 1000 rpm. The other half of the bead suspension was incubated without DNA as a control. After incubation, the beads were washed twice with 16µL incubation buffer and resuspended in 16µL incubation buffer. 2µL of protein and 2µL of radiolabeled prey DNA probes (with a minimum of 5000 counts per minute) were added to both bead suspensions, gently mixed and incubated at 25 ^◦^C for 20 min in an Eppendorf Thermomixer with an Eppendorf Smartblock 1.5 mL at 1000RPM. Incubation buffer, DNA buffer, and protein buffer were designed in such a way to make a constant experimental buffer: 10mM Tris-HCl, pH 7.0, 51mM KCl, 5% v/v glycerol, 1mM spermidine, 1mM DTT, 0.02% Tween20, 1mg mL^−1^ acetylated BSA, 1mM MgCl2. After incubation, the beads were gently washed with 20µL of the same experimental buffer and then resuspended in stop buffer (10mM TrisHCl pH 8.0, 1mM EDTA, 200mM NaCl, 0.2% SDS). The sample was transferred to a liquid Cherenkov-scintillation counter to quantify the radioactive signal. The calculation of DNA recovery (%) was based on a reference sample containing the same amount of labeled 32P 685bp DNA used in each sample. All measurements were performed in triplicate. The data was plotted in Python with the Matplotlib package (see Code availability) [32, 33].

### Ligase-mediated circularization assay

The ligase-mediated circularization assays were performed as previously described [19]. The assays were done with a linear DNA fragment of 240 bp comprising the first 240 bp of the DNA substrate used in the TPM experiments. The 240 bp substrate was generated by PCR amplification of pRD121 [5]. The sequences of the oligonucleotides are available in the Supplementary Information (Supplementary Table 4).

300ng DNA was mixed with HTkC or HTkC R45A at the indicated protein/DNA weight ratios in 50mM Tris, pH 7.0, 75mM KCl. Following incubation at room temperature for 30min, MgCl_2_, ATP, DTT, and T4 DNA ligase were added to final concentrations of 10mM, 1mM, 10mM, and 0.2U*/*µL, respectively, in a total volume of 100µL and incubated for 24h at room temperature. The DNA was purified by phenol/chloroform extraction and ethanol precipitation. Half of the purified DNA was subjected to exonuclease V (New England BioLabs) treatment in CutSmartTM buffer (New England BioLabs), supplemented with ATP to a final concentration of 0.1mM, for 20min at 37 ^◦^C. The exonuclease V was denatured at 70 ^◦^C for 20min. DNA samples were separated on a 2% TAE agarose gel. Gels were stained with 1x GelRed (Biotium) and imaged using the InGenius 5 imaging system (Syngene).

### Tethered particle motion (TPM) experiments

TPM experiments were performed as previously described [35, 36] using 50 mM Tris, pH 7.0, 75 mM KCl as buffer. A standard deviation cutoff of 8% and an anisotropic ratio cutoff of 1.3 were used to select single-tethered beads at all concentrations except 3000 nM HTkC. For 3000 nM HTkC, a standard deviation cutoff of 12% and an anisotropic ratio cutoff of 1.8 were used instead. Measurements at each concentration were done in triplicate. The end-to-end distance for each DNA tether was calculated as 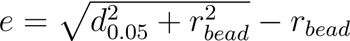 with *r_bead_* being the radius of the polystyrene bead and *d*_0.05_ the top 5% of distances furthest away from the midpoint. Means and standard deviations of the individual measurement series for each concentration were calculated by maximum likelihood estimation assuming a normal distribution. Outliers with a robust Z-score ≥3 or ≤3 were not considered for fitting. The “line to guide the eye” was generated by fitting the means to a logistic function: 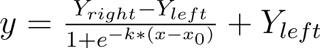. For HTkC, a double logistic function was used instead: 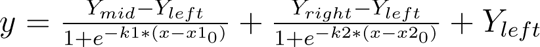. The means of the three individual measurements were averaged for each measured concentration and the standard deviations were error-propagated: 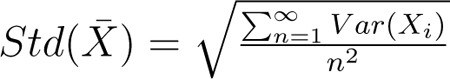 The data was plotted in Python with the Matplotlib package (see Code availability) [32, 33].

### Micrococcal nuclease (MNase) digestion assay

The MNase assays were performed as previously described [19]. The assays were done with the TPM DNA fragment of 685 bp. The 685 bp substrate was generated by PCR amplification of pRD121 [5]. The sequences of the oligonucleotides are available in the Supplementary Information (Supplementary Table 4).

900ng DNA was mixed with HTkC or HTkC R45A at the indicated protein/DNA weight ratios in 50mM Tris, pH 7.0, 75mM KCl. Following incubation at room temperature for 30min, 4U of MNase (New England BioLabs) was added and incubation was continued in digestion buffer, provided by the manufacturer, for 12min at 37 ^◦^C in a total reaction volume of 80µL. The reaction was stopped with EDTA and SDS at a final concentration of 95 mM and 0.5%, respectively. The protected DNA was recovered by phenol/chloroform extraction, followed by ethanol precipitation. The purified DNA was dissolved in 5µL nuclease-free water and separated on a 10% polyacrylamide (0.5x TBE) gel at 120V for 1h with 0.5x TBE as the running buffer. To visualize the DNA, the gel was stained with 1x GelRed (Biotium) and imaged using the InGenius 5 imaging system (Syngene).

### Crystallization, data collection, and structure determination

The complementary oligonucleotides 30 bp-GC50-F and 30 bp-GC50-R (Supplementary Table 4) were annealed at a 1:1 molar ratio to generate the 30 bp-GC50 double-stranded DNA (dsDNA) fragment. For co-crystallization, purified HTkC and dsDNA were mixed in equal volumes, incubated for 30min at 25 ^◦^C, and centrifuged to remove aggregates.

Crystallization trials were carried out in 96-well sitting-drop vapor-diffusion plates using commercial screens, a reservoir volume of 50µL, and drops consisting of 300nL of protein-DNA mixture and 300nL of reservoir solution. The best diffracting crystal was obtained from a drop containing 800µM protein and 200µM dsDNA, and a reservoir solution comprising 0.1M Bis-Tris, pH 6.5, 0.2M MgCl2, and 25% (w/v) PEG 3350. For cryo-protection, crystals were transferred to a droplet of reservoir solution supplemented with 15% (v/v) PEG 400 prior to loop mounting and flash-cooling in liquid nitrogen.

X-ray diffraction data were collected at beamline ID23-1 of the European Synchrotron Radiation Facility (ESRF, Grenoble, France) at 100K, with a wavelength of 0.8856Å, using an EIGER X 16M hybrid pixel detector (Dectris Ltd., Switzerland). Data were indexed, integrated, and scaled in space group C2221 to a resolution of 1.95Å using XDS [37].

The structure was solved by molecular replacement using MOLREP, with the DNA-free HTkC structure (PDB: 9F2C [18]) as the search model, locating one HTkC dimer in the asymmetric unit (ASU) [38]. Following rigid-body refinement with REFMAC5 [39], continuous electron density consistent with dsDNA became apparent throughout the crystal lattice, with 29 nucleotides belonging to the ASU. As the density did not allow unambiguous base assignment, and repeats of 29 nucleotides cannot be explained by the 30-bp dsDNA used for crystallization, the continuous DNA density is likely the result of a stochastic distribution of the short duplex fragments along a common DNA trajectory in the crystal lattice. Consequently, we decided to model this density as a continuous 29-nt DNA in the ASU with the sequence of primer 30 bp-GC50-F, even though this resulted in imperfect base pairing in the symmetry-generated dsDNA and small discontinuities in the seemingly endless dsDNA. The structure was finalized through iterative rounds of manual modeling in Coot and refinement in REFMAC5 [39, 40].

Data processing and refinement statistics are summarized in Supplementary Table 3, and coordinates and structure factors are available in the Protein Data Bank under accession code 9T30.

### Molecular dynamics (MD) simulations

For MD simulations, we used GROMACS [41] with the AMBER ff14sb-ParmBSC1 force field [42, 43]. All systems were solvated in a dodecahedron box with a distance of at least 1.0 nm to the box boundary and with the TIP3P water model [44]. Water molecules were replaced at random with Na+ and Cl-ions to charge-neutralize the system. 75 mM of Na+ and Cl-ions were additionally added. Energy minimization was performed using the steepest descent method for 5000,000 steps until the largest force was below 1000.0 kJ/mol/nm. To equilibrate the solvent and ions, heavy atoms were position constrained for 100 ps at 298 K and 1 bar. The cutoff for van der Waals interactions was 1.1 nm. Electrostatic interactions beyond a cutoff of 1.1 nm were treated with the particle-mesh Ewald method using a grid spacing of 0.16 nm. Temperature and pressure were kept constant with the Vrescale thermostat (Bussi-Donadio-Parrinello thermostat) [45] and the Parrinello-Rahman barostat [46], respectively. Bonds were constrained with LINCS [47] and simulations were conducted in time steps of 2 fs.

The starting structure for the MD, with DNA bent around the HTkC tetramer, was constructed from co-crystal structure (PDB: 9T30) (Supplementary Fig. 5a). One tetramer with 75 base pairs of DNA was isolated from the crystal lattice. The separate DNA molecules were linked together and the sequence of one strand was altered to create functional base pairing between both strands. Missing N-terminal residues were added from a tetrameric AlphaFold2 prediction of HTkC [18] and any atoms that were missing on residues were filled in. For HMfB, the PDB structure 5T5K was used as the starting structure [4]. The two single-stranded nucleotides at the start and end of the DNA were removed.

The minor and major groove widths and the atoms involved in protein-DNA contacts were identified as described in a previous study by van Heesch et al. [48]. The bending angle of the DNA was computed by first converting the DNA base pairs into a rigid body model [49], which defines mean reference frames for each base pair. Then, after defining vectors through the origins of the mean reference frames of the G13-C63 base pair and T44-A32 (center) base pair, and the A75-T1 base pair and T44-A32 (center) base pair, the bending angle was computed as 180^◦^ minus the inverse cosine of the dot product of these vectors: 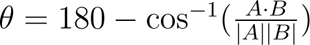. The MD data was plotted in Python with the Matplotlib package (see Code availability) [32, 33].

### Deletion of *htkC* from *Thermococcus kodakarensis*, RNAseq, and growth curve measurements

Marker-less deletion of *htkC* (TK1040) in *T. kodakarensis* was performed using a pop-in/pop-out approach [50]. This method uses the *T. kodakarensis* strain KU216 as a parental strain, which lacks the orotidine-5’-monophosphate decarboxylase gene *pyrF*. The Δ*pyrF* mutation confers uracil auxotrophy as well as the resistance to 5-fluoroorotic acid (5-FOA). To construct a deletion plasmid targeting TK1040, two ∼1-kb regions immediately upstream and downstream of the coding sequence were PCR amplified using genomic DNA of *T. kodakarensis* KU216 as a template. The sequences of the oligonucleotides are available in the Supplementary Information (Supplementary Table 4).

The two fragments were cloned together into the BamHI-EcoRI site of the pUD3 vector, containing the *pyrF* marker [51], using In-Fusion HD Cloning Plus reagents (Takara Clontech, 638910). The generated deletion plasmid was used for transformation of KU216 to integrate the construct into the target genomic region. Transformants with the marker were selected in the uracil-free medium ASW-AAm1-S^0^ [52]. To select strains in which pop-out of the *pyrF* marker occurred, cells were further grown in solid ASW-YT-m1-S^0^ medium [53] supplemented with 8.3 g/L of 5-FOA•H2O and 44 mM NaOH. Obtained 5-FOA resistant cells were analyzed by PCR to confirm the marker-less deletion of *htkC*. The Δ*htkC* mutant and the parental strain KU216 were grown in ASW-YT-m1-S^0^ and subjected to growth curve measurement and RNA-seq as described previously [29]. RNA-seq was performed in three biological replicates using cells collected at an OD660 of ∼0.15. The RNA-seq and growth curve data were plotted in Python with the Matplotlib package (see Code availability) [32, 33].

## Data availability

The TPM data, raw gel images, MD data, analysed RNA-seq data, growth curve data, CD spectroscopy data, SEC-MALS data, MD trajectories, and MD movies are available at 4TU with the DOI 10.4121/fa8bd0b5-a15d-496c-9a03-86fe1c38c2ee. X-ray structural data have been deposited in the PDB database with accession number 9T30.

## Code availability

The source code for the analysis of data and generation of figures is available at 4TU with the DOI 10.4121/fa8bd0b5-a15d-496c-9a03-86fe1c38c2ee.

## Acknowledgments

This research was funded by the Dutch Research Council [OCENW.GROOT.2019.012] to R.T.D and by institutional funds from the Max Planck Society.

We are grateful to the staff of Beamline ID23-1 of the European Synchrotron Radiation Facility (ESRF, Grenoble, France) for excellent technical support. We extend our thanks to Reinhard Albrecht for assistance with crystallization and crystallographic data collection. V.A. and B.H.A. would like to thank Andrei Lupas for his continued support.

We would like to thank Aimee L. Boyle for thorough reading and reviewing of the manuscript. The ALICE HPC cluster at Leiden University is kindly acknowledged for providing the infrastructure necessary to perform many of the computations described in this article.

UCSF ChimeraX is kindly acknowledged for the molecular graphics and analyses in this article.

## Author contributions

**Samuel Schwab:** Writing - original draft, Conceptualization, Visualization, Investigation, Methodology, Formal analysis

**Yimin Hu:** Writing - review & editing, Investigation, Visualization, Methodology

Shingo Yamamoto: Writing - review & editing, Investigation

Kodai Yamaura: Writing - review & editing, Investigation

Marc K. Cajili: Writing - review & editing, Investigation

Bert van Erp: Writing - review & editing, Investigation

**Naomichi Takemata:** Writing - review & editing, Investigation, Methodology, Formal analysis

Haruyuki Atomi: Writing - review & editing, Supervision

Marcus D. Hartmann: Writing - review & editing, Investigation, Methodology

Vikram Alva: Writing - review & editing, Supervision

Birte Hernandez Alvarez: Writing - review & editing, Supervision

Remus T. Dame: Writing - review & editing, Supervision, Funding acquisition, Conceptualization

## Declaration of interests

The authors declare no conflict of interests.

## Notes

### Competing Interest Statement

The authors have declared no competing interest.

