## Supplementary Info for "Archaeal histone HTkC hypercompacts DNA"

### Supplementary Information

Supplementary Table 1: Theoretical and measured (SEC-MALS) molecular weights.

| Protein | Theoretical MW of monomer (Da) | SEC-MALS measured MW (Da) |
|---|---|---|
| HTkC | 6752 | $2.708 * 10^4 \pm 1.253 * 10^2$ |
| HTkC R45A | 6667 | $1.316 * 10^4 \pm 9.265 * 10^1$ |

Supplementary Table 2: Plasmids created for this study.

| Name | Backbone | Insert | Resistance | Addgene # |
|---|---|---|---|---|
| pRD661 | pET30b | HTkC R45A | Kanamycin | XXX |

Supplementary Table 3: Data collection and refinement statistics for HTkC-DNA. Values for the outer shell are given in parentheses.

| | **HTkC-DNA** |
|---|---|
| **Data Collection** | |
| Space group | $C222_1$ |
| Cell dimensions | |
| $a, b, c$ (Å) | 63.22, 80.90, 72.64 |
| $\alpha, \beta, \gamma$ (°) | 90, 90, 90 |
| Resolution range (Å) | 41.12-1.95 (2.07-1.95) |
| Completeness (%) | 95.4 (93.2) |
| Redundancy | 7.57 (7.85) |
| $< \frac{I}{\sigma(I)} >$ | 11.99 (2.14) |
| $R_{meas}$ | 0.093 (0.849) |
| **Refinement** | |
| No. of reflections, working set | 12636 |
| No. of reflections, test set | 666 |
| Final $R_{cryst}$ | 0.226 |
| Final $R_{free}$ | 0.278 |
| R.m.s. deviations | |
| Bonds (Å) | 0.0038 |
| Angles (°) | 1.055 |

Supplementary Table 4: Oligonucleotides used in this study.

| Name primer | Used for | Sequence (5'-3') |
|---|---|---|
| HTkC R45A Ins Fwd | pRD661 | CAGGGTTTTACGACCTTCTGCCTGTGCCGCTT TAATTGCT |
| HTkC R45A Ins Rev | pRD661 | TTTAAGAAGGAGATATACATATGGCAGAAATG CTGGTTAA |
| HTkC R45A Vec Fwd | pRD661 | TTAACCAGCATTTCTGCCATATGTATATCTCC TTCTTAAAGTTAAACA |
| HTkC R45A Vec Rev | pRD661 | AGCAATTAAAGCGGCACAGGCAGAAGGTCGTA AAACCCTG |
| 150-bp Fwd | EMSA | TTACTTTCACCAGCGTTTCTGGGTGAGCAAAA ACAG |
| 150-bp Rev | EMSA | CGCTCATGAGACAATAACCCTGATAAATGCTT CAATAATATTG |
| 240-bp Fwd [Phos] | Circ. | [Phos]TTACTTTCACCAGCGTTTCTGGGTGA GCAAAAACAG |
| 240-bp Rev [Phos] | Circ. | [Phos]TGGTTTCTTAGACGTCAGGTGGCACT TTTCGG |
| 685-bp Fwd [Biotin] | TPM | [Biotin]TTACTTTCACCAGCGTTTCTGGGT GAGCAAAAACAG |
| 685-bp Rev [DIG] | TPM | [DIG]CCAAGTAGCGAAGCGAGCAGGACTGGG CGG |
| 685-bp Fwd | MNase | TTACTTTCACCAGCGTTTCTGGGTGAGCAAAA ACAG |
| 685-bp Rev | MNase | CCAAGTAGCGAAGCGAGCAGGACTGGGCGG |
| 1-kb upstream Fwd | Δ*htkC* | CAGGTCGACTCTAGAGGATCCTTCGGCTCTCT TCTCAGGGT |
| 1-kb upstream Rev | Δ*htkC* | GGCAGAAAATGGCTAACACCTCCTTATTGGCC T |
| 1-kb downstream Fwd | Δ*htkC* | GGTGTTAGCCATTTTCTGCCATCTTTCAGTTT TTCC |
| 1-kb downstream Rev | Δ*htkC* | TATGACCATGATTACGAATTCAGGAAGCTCCG TTGAACTCG |
| 30-bp GC50 Fwd | Crystal | TTAAAGCCCGTTAAAGCCCGTTAAAGCCCG |
| 30-bp GC50 Rev | Crystal | CGGGCTTTAACGGGCTTTAACGGGCTTTAA |

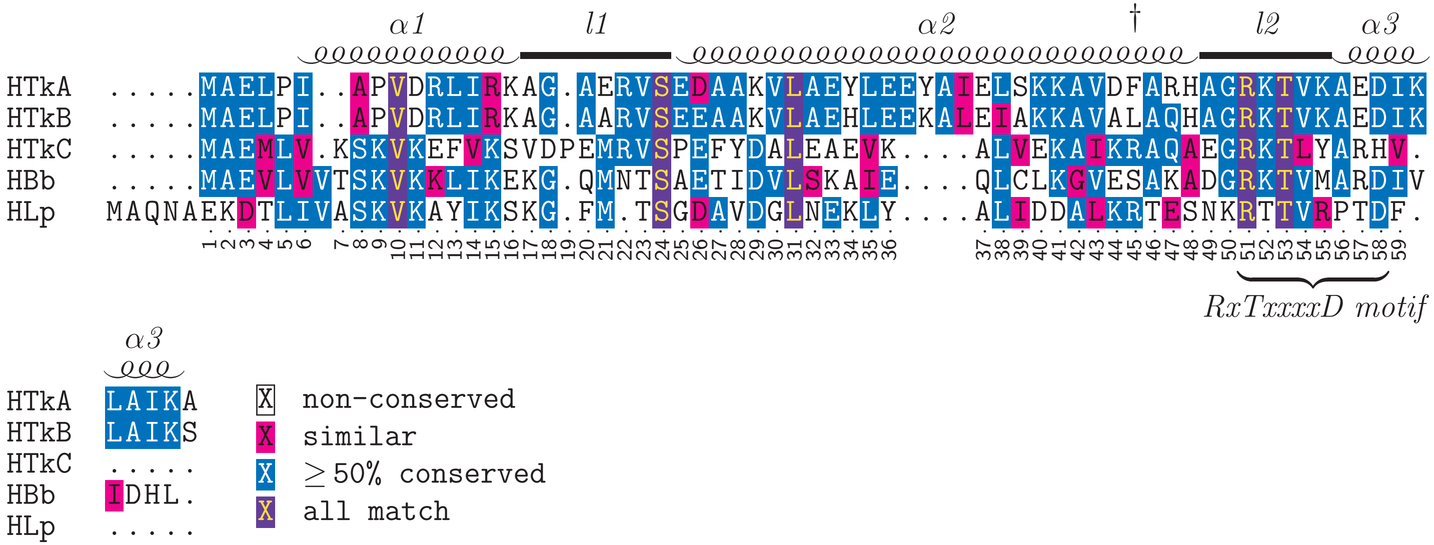

Supplementary Figure 1: **Multiple sequence alignment of hypernucleosome-forming histones HTkA and HTkB, face-to-face histones HTkC and HLp, and bacterial dimer histone HBb.** Helices $\alpha1$, $\alpha2$, and $\alpha3$ and loops l1 and l2 are annotated above the sequences. The dagger (†) marks arginine 45. The ruler below the sequences shows HTkC's amino acid numbering. Alignment was made with MUSCLE5 using default settings [1].

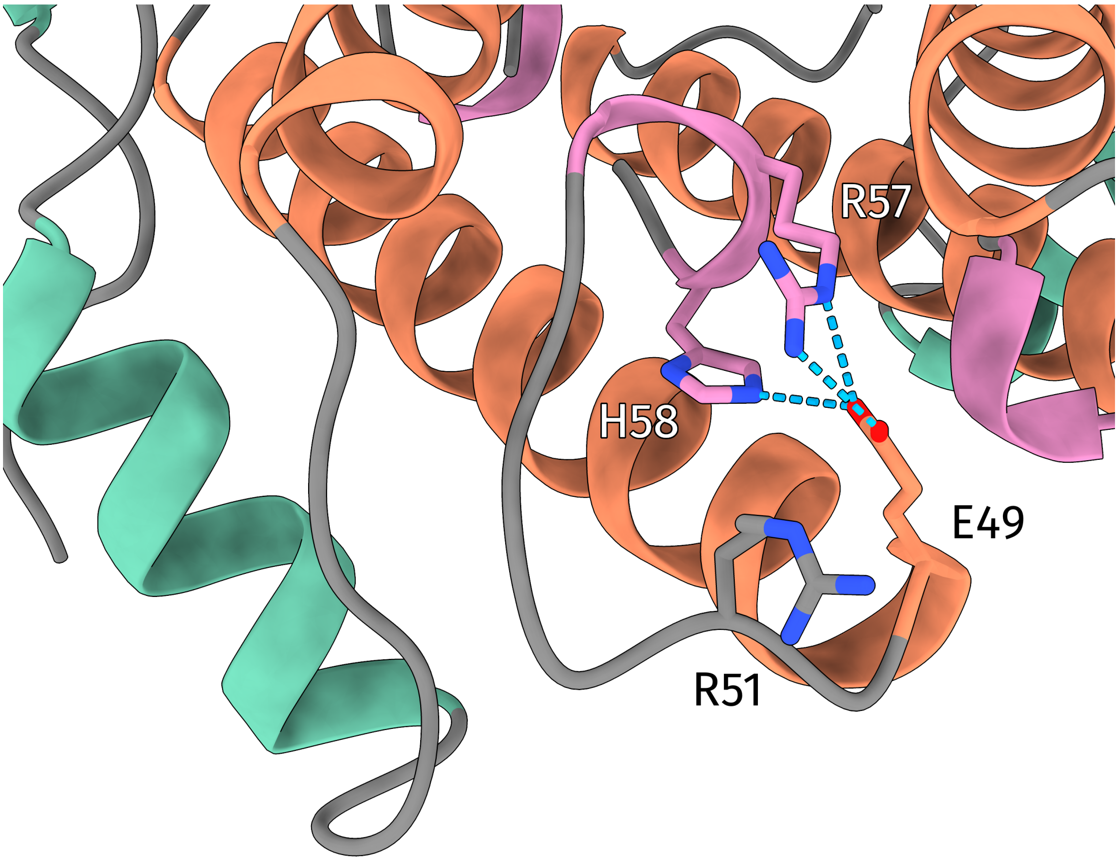

Supplementary Figure 2: **Close-up of the l2 loop of HTkC (PDB: 9T30).** The l2 loop and helix $\alpha3$ are positioned by E49, R57, and H58. R51, which in other histones positions the l2 loop and helix $\alpha3$, faces outwards.

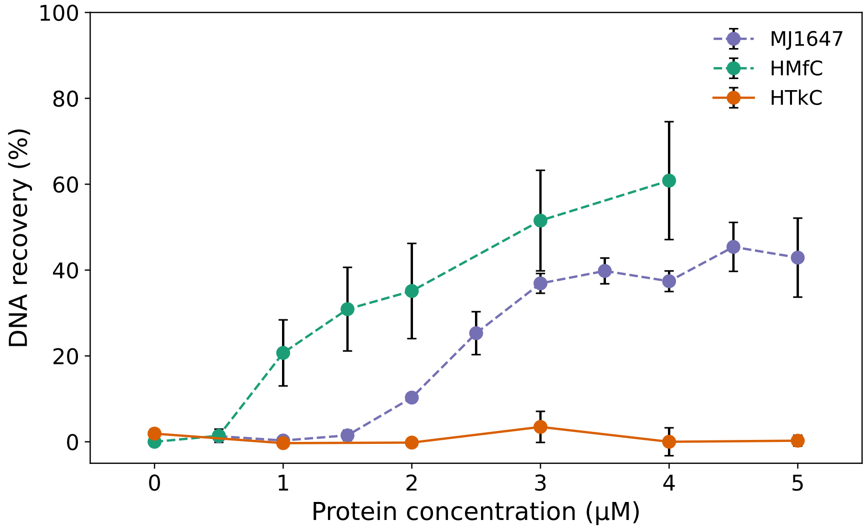

(a)

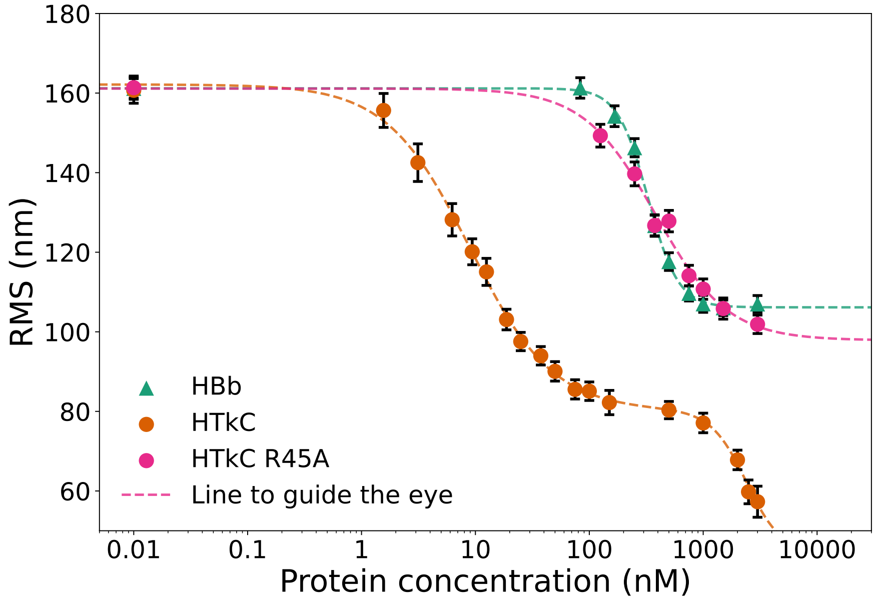

(b)

Supplementary Figure 3: **Supplementary DNA-bridging and TPM data.** (a) DNA bridging assay with HTkC. DNA-bridging histones HMfC and MJ1647 are shown as references [2, 3]. DNA bridging activity is represented on the y-axis as the percentage of radioactively labeled DNA recovered after pulling-down the magnetic-bead immobilized DNA. Points represent the mean values of three independent measurements. One standard deviation is visualized as error bars. (b) The RMS values from TPM of HTkC, HTkC R45A, and HBb in orange, pink, and green respectively.

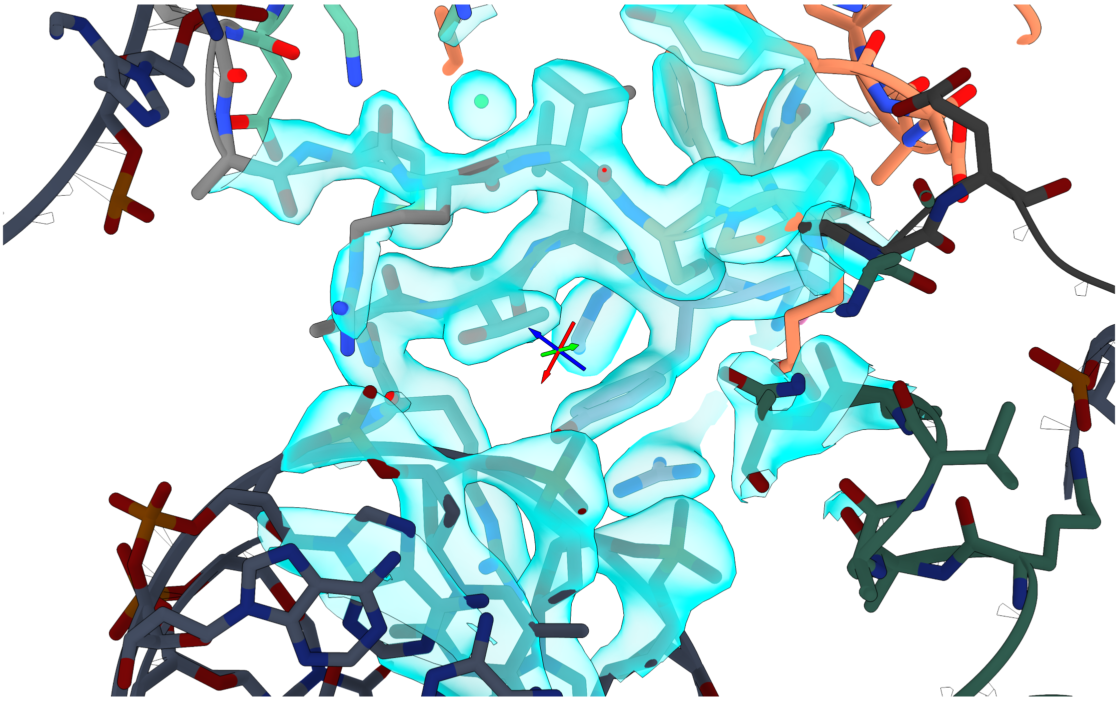

Supplementary Figure 4: **Density map of the HTkC-DNA crystal.** 2mFo-DFc map of HTkC's l1 and l2 loop region contoured at 1.5 sigma (cyan).

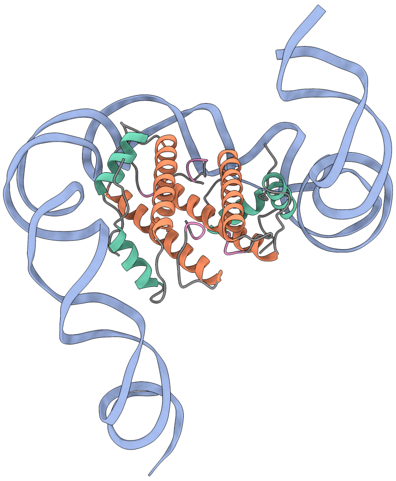

(a)

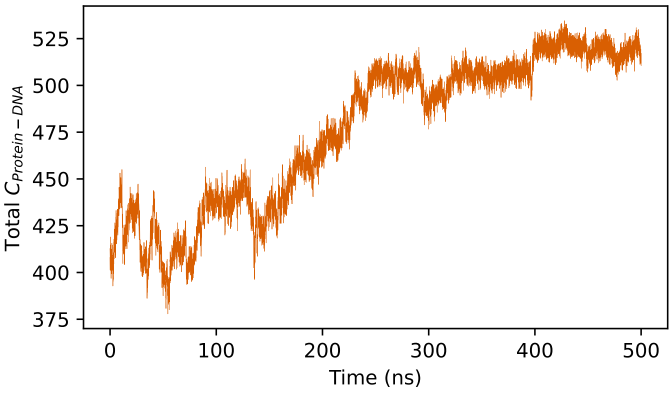

(b)

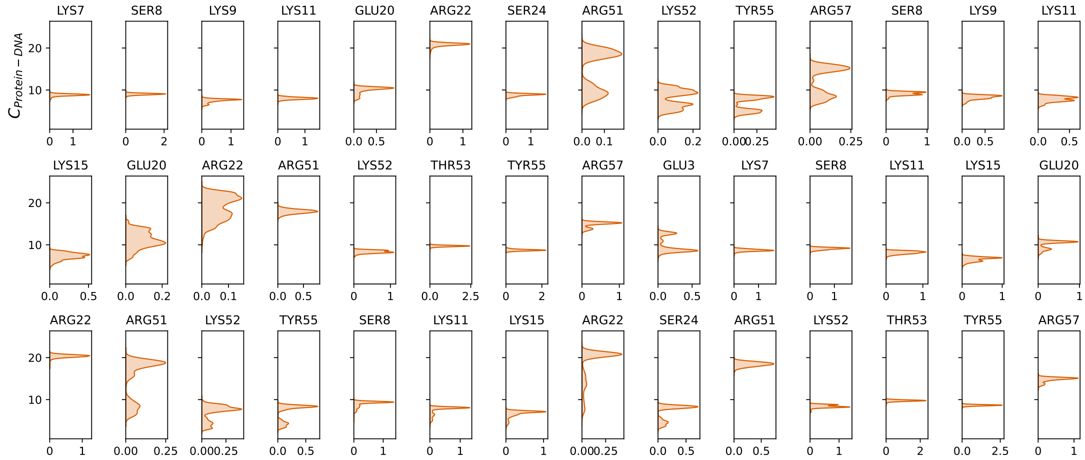

(c)

Supplementary Figure 5: (Figure continues on the next page.)

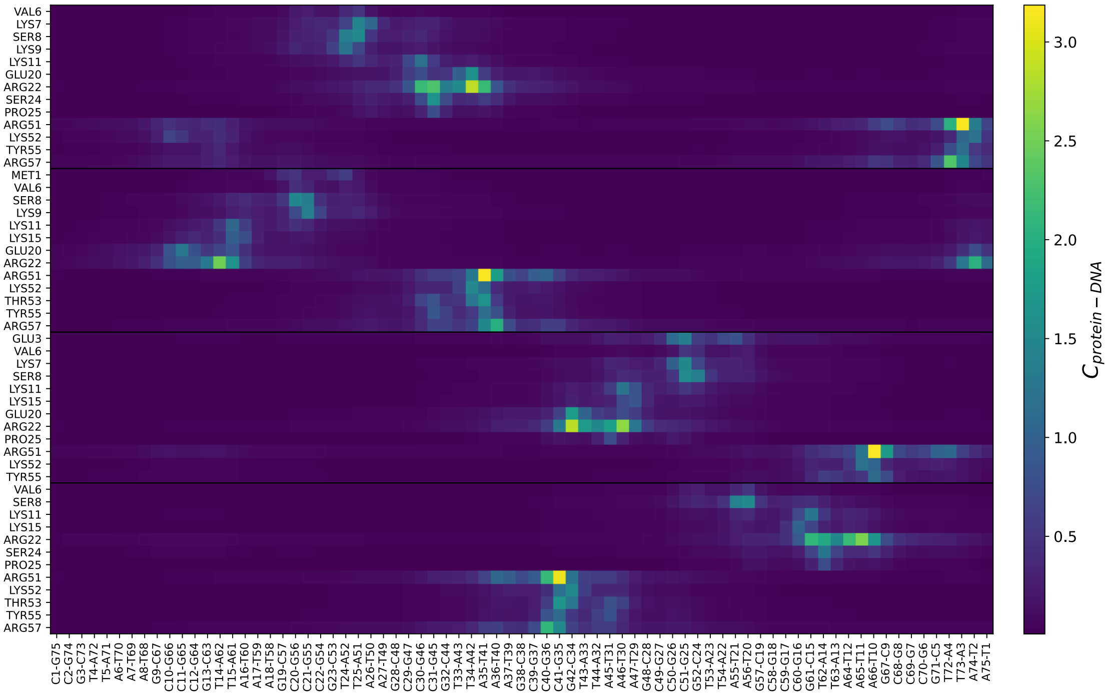

(d)

Supplementary Figure 5: **MD simulation #1** (a) Starting structure for both MD runs. (b) Total number of protein-DNA contacts during simulation #1. (c) Decomposed density estimates of the number of contacts for the fully wrapped state (the last 250 ns) of each protein residue with respect to DNA of simulation #1. Only residues with an average number of contacts above 5 are shown. The data for each monomer are concatenated. (d) Heat map of the mean number of contacts for the fully wrapped state (the last 250 ns) of each protein residue with each base pair for simulation #1. The data for each protein monomer is concatenated with a black dividing line separating the data of each monomer.

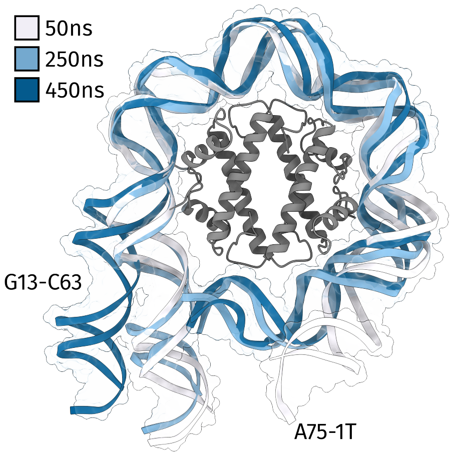

(a)

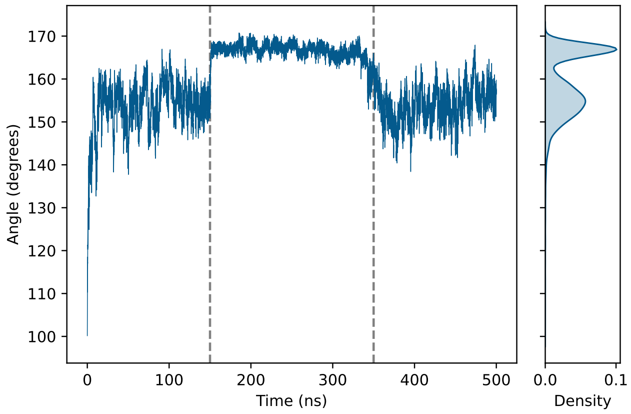

(b)

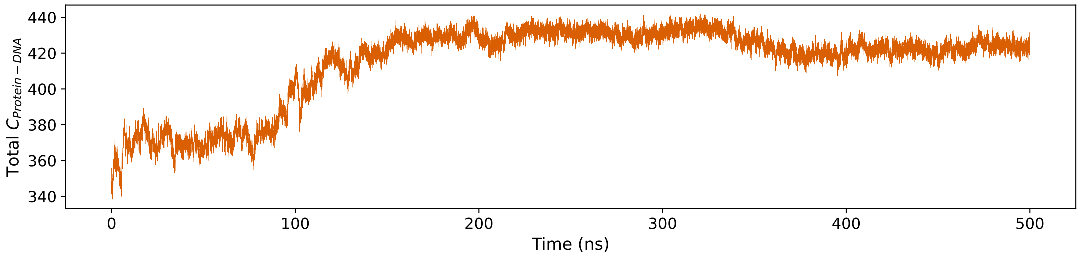

(c)

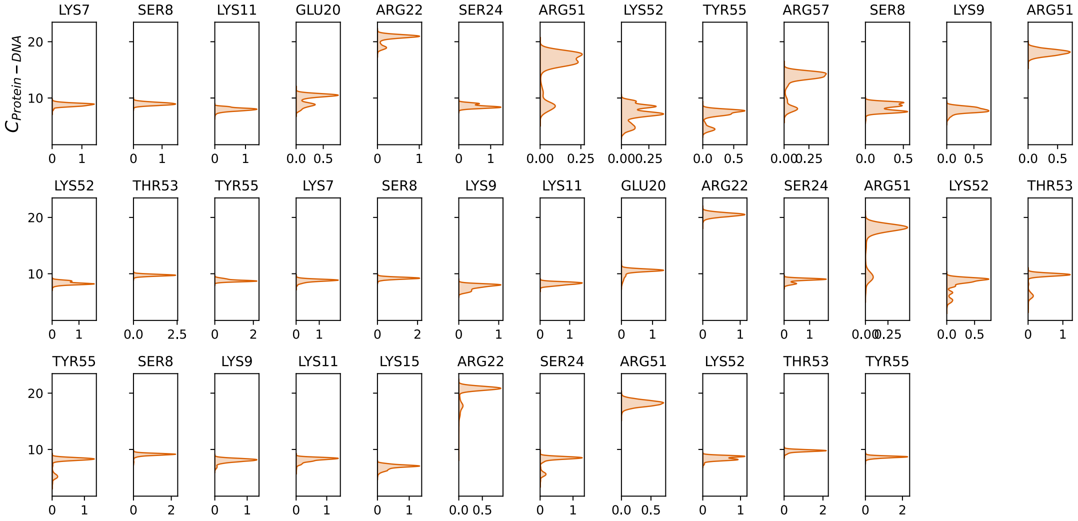

(d)

Supplementary Figure 6: (Figure continues on the next page.)

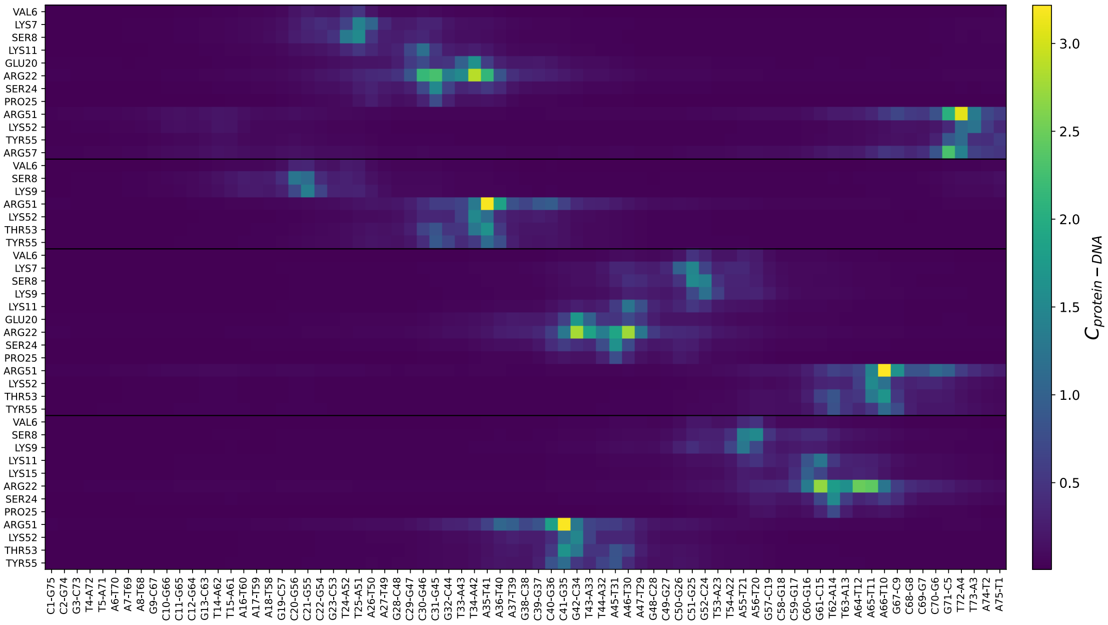

(e)

Supplementary Figure 6: **MD simulation #2** (a) Three frames, at 50, 250, and 450 ns, from simulation #2, that show the trajectory of wrapping and unwrapping. (b) The bending angle in degrees between base pairs G13-C63, A75-T1, and the middle base pair (C30-G46) during simulation #2. The time points where complete wrapping occurs (150 ns) and where slight unwrapping occurs (350 ns) are highlighted by the gray dashed line. (c) Total number of protein-DNA contacts during simulation #2. (d) Decomposed density estimates of the number of contacts for the fully wrapped state (150 ns-350 ns) of each protein residue with respect to the DNA of simulation #2. Only residues with an average number of contacts above 5 are shown. The data for each monomer are concatenated. (e) Heat map of the mean number of contacts for the fully wrapped state (150 ns-350 ns) of each protein residue with each base pair for simulation #2. The data for each protein monomer is concatenated with a black dividing line separating the data of each monomer.

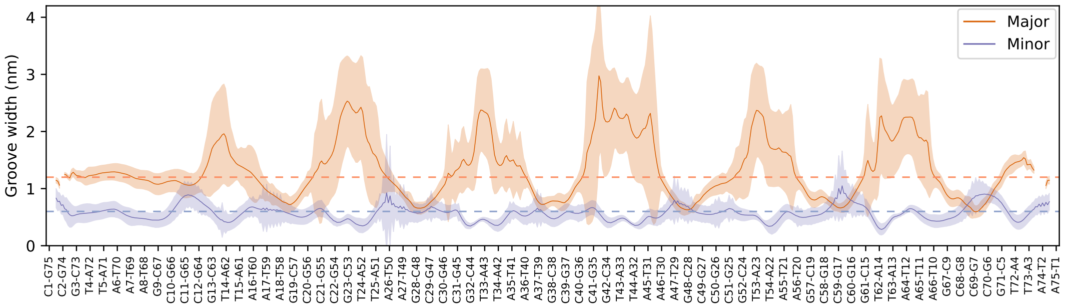

(a)

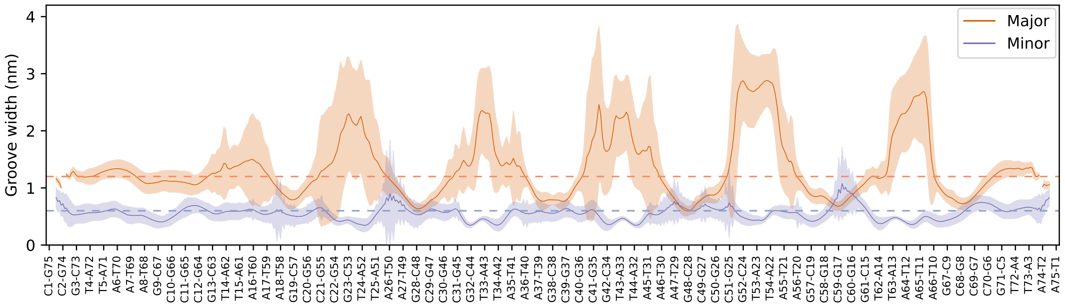

(b)

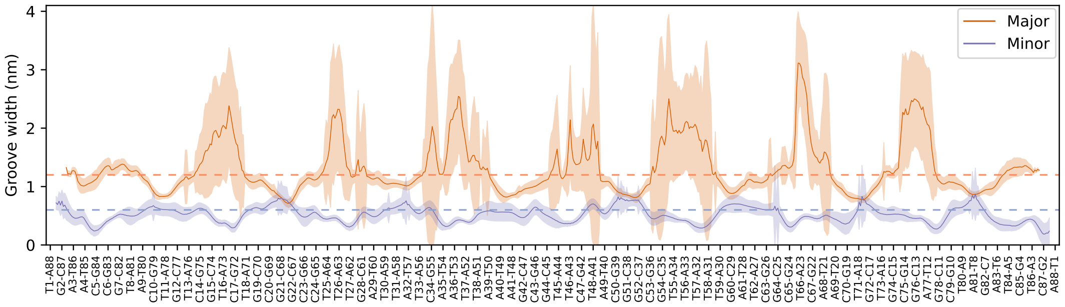

(c)

Supplementary Figure 7: **Major and minor grooves during the MD simulations.** (a,b) Major and minor groove widths, in orange and blue respectively, of the HTkC-bound DNA molecules of simulation #1 (a) and #2 (b) during their fully wrapped states. The shaded area represents one standard deviation. The normal groove width of B-DNA is indicated by the striped line. (c) Major and minor groove widths of a 1 ns simulation of the HMfB hypernucleosome crystal structure (PDB: 5T5K).

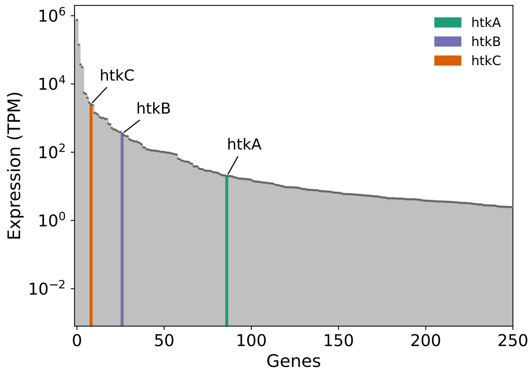

(a)

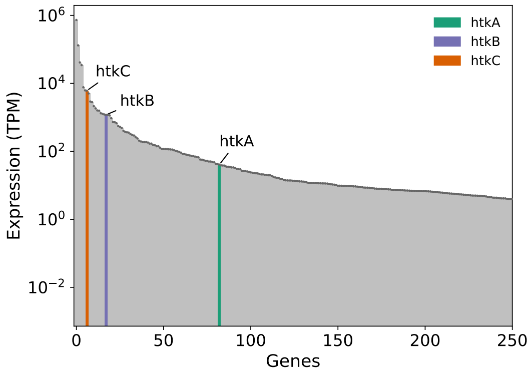

(b)

Supplementary Figure 8: **Expression data from biological replicates 2 and 3.** (a,b) Expression levels, calculated as Transcripts per Kilobase Million (TPM), of the top 250 expressed genes from biological replicate 2 (a) and 3 (b). Genes are ordered from highest (left) to lowest (right) expression. The *htkA*, *htkB*, and *htkC* genes are highlighted in green, purple, and orange respectively.
